# How the olfactory bulb maintains stable odor manifolds amid adaptation and representational drift

**DOI:** 10.64898/2026.01.23.701335

**Authors:** CiCi Xingyu Zheng, Bin Yu, Saket Navlakha, Alexei Koulakov

## Abstract

Adaptive coding in sensory circuits enables stable perception while accommodating experience-dependent changes. In the olfactory bulb (OB), repeated odor exposure reshapes population activity even without explicit behavioral feedback, but the underlying circuit mechanisms remain unclear. By analyzing longitudinal two-photon calcium imaging datasets from the mouse OB, we identified three concurrent forms of representational change: gain adaptation, similarity-dependent pattern separation or convergence, and a rotation of encoding subspace resulting in the representational drift. Using a computational model of the mitral cell-granule cell circuit, we showed that Hebbian plasticity and structural connectivity constraints are sufficient to reproduce these transformations. Despite global representational drift, the relative geometry of odor response vectors remained stable, preserving a low-dimensional odor manifold. Together, our results reveal how local plasticity and network structure jointly enable both stability and flexibility in early sensory coding.

## Introduction

Adaptive coding allows sensory systems to extract behaviorally relevant information from continuously changing environments. In olfaction, adaptation occurs across multiple timescales from rapid habituation within a few breaths (Chaudhury et al., 2010; Kim et al., 2020; Shen et al., 2020) to long-term adjustments spanning days or even seasons (Hamdani et al., 2008; Teşileanu et al., 2019). Adaptation can decorrelate sensory representations, facilitating flexible behaviors such as distinguishing food sources from predators or compensating for persistent background odors (Dalton, 2000). Adaptive coding is not a single process: it can take distinct forms across multiple levels, from receptor and synaptic plasticity to circuit-level reorganizations. Understanding how odor representations evolve over days, and what computational consequences these changes entail, is therefore essential for linking sensory experience to long-term learning.

The mammalian olfactory system transforms chemical stimuli from odorants into structured neural representations through a series of processing stages. Odorants are detected by olfactory sensory neurons (OSNs) in the nasal epithelium, each expressing one odorant receptor (OR) gene out of hundreds of possibilities (Buck & Axel, 1991; Chess et al., 1994; Malnic et al., 1999). OSNs expressing the same receptor send their axons to a small number of glomeruli located in the glomerular layer of the olfactory bulb (OB) (Mombaerts et al., 1996). The glomeruli form a dense two-dimensional sheet-like structure across the OB surface, creating a stereotypical spatial map of inputs from the olfactory epithelium (Bozza et al., 2009; Giessel & Datta, 2014; Wang et al., 1998). Within each glomerulus, OSNs expressing a specific OR gene type form excitatory synapses onto the principal OB projection neurons, mitral and tufted cells (MCs and TCs), as well as onto local interneurons (Mombaerts, 2006).

Before transmitting information to cortical targets such as the piriform cortex, MCs and TCs interact with inhibitory interneurons in the deeper granule cell (GC) layer of the OB (Egger & Urban, 2006). GCs form reciprocal dendrodendritic synapses with MC/TCs, where excitation and inhibition occur at the same contact site (Aghvami et al., 2022; Aroniadou-Anderjaska et al., 1999; Rall et al., 1966). These reciprocal connections mediate recurrent inhibition that may dynamically regulate overall network gain (Shepherd et al., 2007; Yokoi et al., 1995). Because GCs integrate inputs from MC/TCs associated with multiple glomeruli, they implement a powerful form of lateral inhibition that can reshape the principal neurons’ activity based on combinatorial odor inputs (Arevian et al., 2007; Cleland & Linster, 2012; Koulakov & Rinberg, 2011). MCs in particular, exhibit more concentration-invariant responses than TCs and provide the dominant projection to piriform cortex, making them a good substrate for studying long-term changes in population coding (Chae et al., 2022). MCs and TCs subsequently convey odor information to the higher brain regions, including piriform cortex (PCx) and anterior olfactory nucleus (AON), for further processing (Shepherd, 2003).

Circuits within these cortical regions are known to support odor learning, associative processing, and integration with other sensory and contextual information (Boyd et al., 2012; Calu et al., 2007; Large et al., 2016; Otazu et al., 2015) via long-term changes in olfactory representations (Li et al., 2006; Wilson, 2000; Wilson et al., 2006). Numerous studies have shown that plasticity in the PCx recurrent connectivity supports perceptual and discriminative learning, as well as reward-associated memory formation (Bao et al., 2016; Cohen et al., 2008; Franks et al., 2011; Gadziola et al., 2020). At the same time, population activity in PCx has been reported to drift over days, even during infrequent or passive odor exposure (Schoonover et al., 2021). It remains unclear, however, how much of this representational change arises within the cortex itself versus being inherited from earlier processing stages. One step upstream, the OB already exhibits time and exposure-dependent adaptations in its output patterns (Chu et al., 2016; Kato et al., 2012). Since recordings from OSNs have shown stable synaptic release properties during passive odor experience (Chu et al., 2017), these longer-timescale changes in neural responses could emerge within the OB networks. Such bulb-level plasticity could thus serve as a source of the representational drift observed in the cortex.

Despite its role as an early sensory relay, OB processes olfactory information in a dynamic and experience-dependent manner (Chu et al., 2016; Wu et al., 2020; Yu et al., 2025). During wakefulness, repeated passive exposure to the same odorants leads to a gradual, odor-specific reduction in MC responses (Chu et al., 2016; Kato et al., 2012; Yamada et al., 2017). Moreover, olfactory perceptual learning, where animals learn to discriminate initially indistinguishable odorants, has been shown to involve changes as early as the bulb, including the decorrelation of MC and TC population activity for similar odors, particularly pronounced in the MCs (Chu et al., 2016; Yamada et al., 2017). These findings indicate that the OB is not merely a stable encoder of sensory inputs but a flexible circuit capable of long-term adaptation.

Such experience-dependent decorrelation can be mediated by inhibitory plasticity that refines local circuits to selectively suppress overlapping activity patterns among MCs. In agreement with this idea, GCs enhance pattern separation during olfactory perceptual learning: optogenetic activation of GCs increases separation of MC population responses and improves odor discrimination performance, whereas GC inactivation reduces separation and impairs discrimination (Alonso et al., 2012; Li et al., 2018). In rodents, OB is also one of the few brain regions where new inhibitory neurons are continually incorporated throughout life, suggesting an additional source of long-term circuit plasticity (Lledo & Valley, 2016). However, adult-born GCs integrate gradually and may contribute primarily to later phases of adaptation (Whitman & Greer, 2007); their ablation does not abolish all forms of learning-related plasticity (Li et al., 2018; Nissant et al., 2009). Together, these findings point to the MC-GC recurrent network as a candidate for shaping odor representations over extended timescales.

Here, we investigate how odor representations by the MCs, one of the OB’s principal output neurons, evolve across days of repeated passive exposure. By analyzing longitudinal two-photon calcium imaging data from mice repeatedly exposed to either similar or dissimilar odor pairs and to an eight-odor mixture panel, we quantified how population codes transform over time. We then developed a recurrent inhibitory feedback model of the MC-GC circuit to test whether local plasticity and circuit structure can account for the observed transformations. Together, our work unifies several distinct forms of representational changes – pattern separation, pattern convergence, gain adaptation, and representational drift – under a single circuit mechanism.

## Results

Our goal is to investigate how odor representations of MCs in the OB change under repeated stimulus exposure over several days. Before proceeding with our analyses, we define four types of long-term plasticity in odor representations: gain adaptation, pattern separation, pattern convergence, and representational drift (Fig. 1). To characterize these changes, we consider the population response vector, whose components correspond to the responses of individual MCs to an odorant. *Gain adaptation* is defined as a reduction in the length of the population response vector (Fig. 1A). *Pattern separation* can occur when pairs of odors are presented intermittently across multiple experimental sessions over several days. It corresponds to a rotation of the population response vectors away from each other while remaining within the same plane (Fig. 1B). *Pattern convergence* is the opposite effect and consists of a rotation of population responses to odor pairs toward each other (Fig. 1C). Finally, *representational drift* is defined as a rotation of the plane containing population responses to the presented odors as a result of odor exposure (Fig. 1D). While the first three phenomena have been reported in the OB (Chu et al., 2016; Kato et al., 2012), representational drift remains unexplored. Below, we combine a computational model of OB circuitry with longitudinal calcium imaging data from mice repeatedly exposed to odors (similar or dissimilar pairs, as well as an eight-odor mixture panel) over 7–9 days (Chu et al., 2016; Li et al., 2018). This integrated approach enables us to identify the circuit features underlying each type of plasticity and to test theoretical predictions against experimental observations.

**Figure 1.**
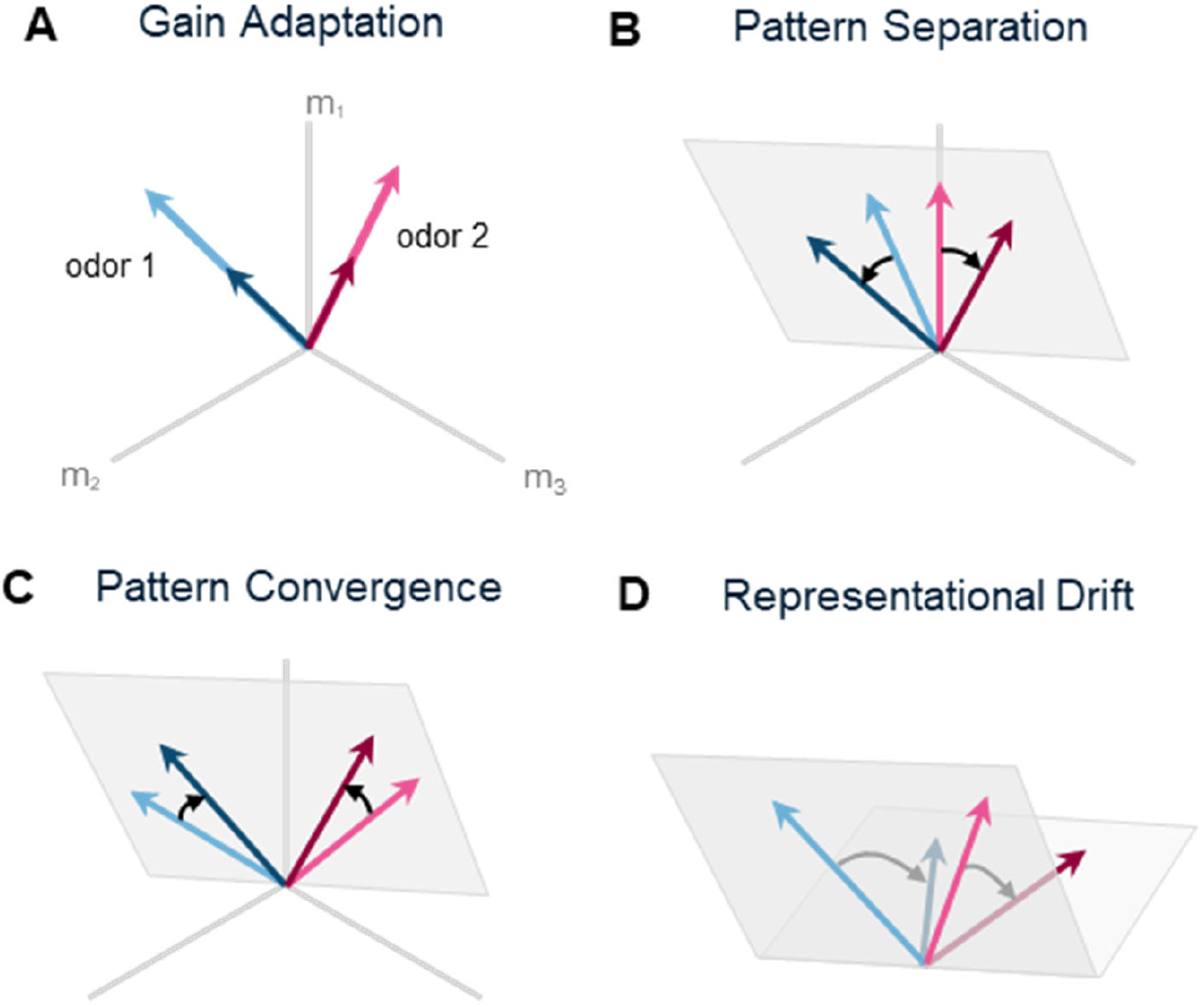
Schematic representation of four types of adaptation of odor representations resulting from odor exposure. (**A**) *Gain adaptation*. Repeated exposure to an odor leads to a reduced response. Light blue and pink arrows represent population responses (m_1_, m_2_, m_3_, … are responses of the individual mitral cells) to odor 1 and odor 2 during early odor exposure; darker arrows show representations of the same odors after extended exposure. (**B**) *Pattern separation*. Initially similar odor representations (light arrows, small angular separation) become more distinct over time, increasing the angle between representation vectors (darker arrows). The vectors are limited to move in the same subspace (shaded). (**C**) *Pattern convergence*. Initially distinct odor representations become more similar over time, decreasing the angle between representation vectors, while remaining in the same subspace (shaded). (**D**) *Representational drift*. Odor representation subspace may rotate over time while maintaining relative relationships between population response vectors. Curved arrows indicate the drift direction.

### Gain adaptation reflects long term population-level activity decay

We define long-term gain adaptation as the gradual reduction in the MC population activity over days of repeated odor exposure. This phenomenon manifests itself as decreased amplitude of MC population responses (Fig. 2A; Fig. S2A-D). We hypothesized that this long-term adaptation arises from activity-dependent plasticity in the dendrodendritic synapses between MCs and GCs (Fig. 2B). Odor-evoked MC activation drives GC responses, which in turn provide inhibitory feedback to MCs. Hebbian strengthening of these reciprocal connections progressively enhances inhibitory feedback, leading to reduced MC responses over time.

**Figure 2.**
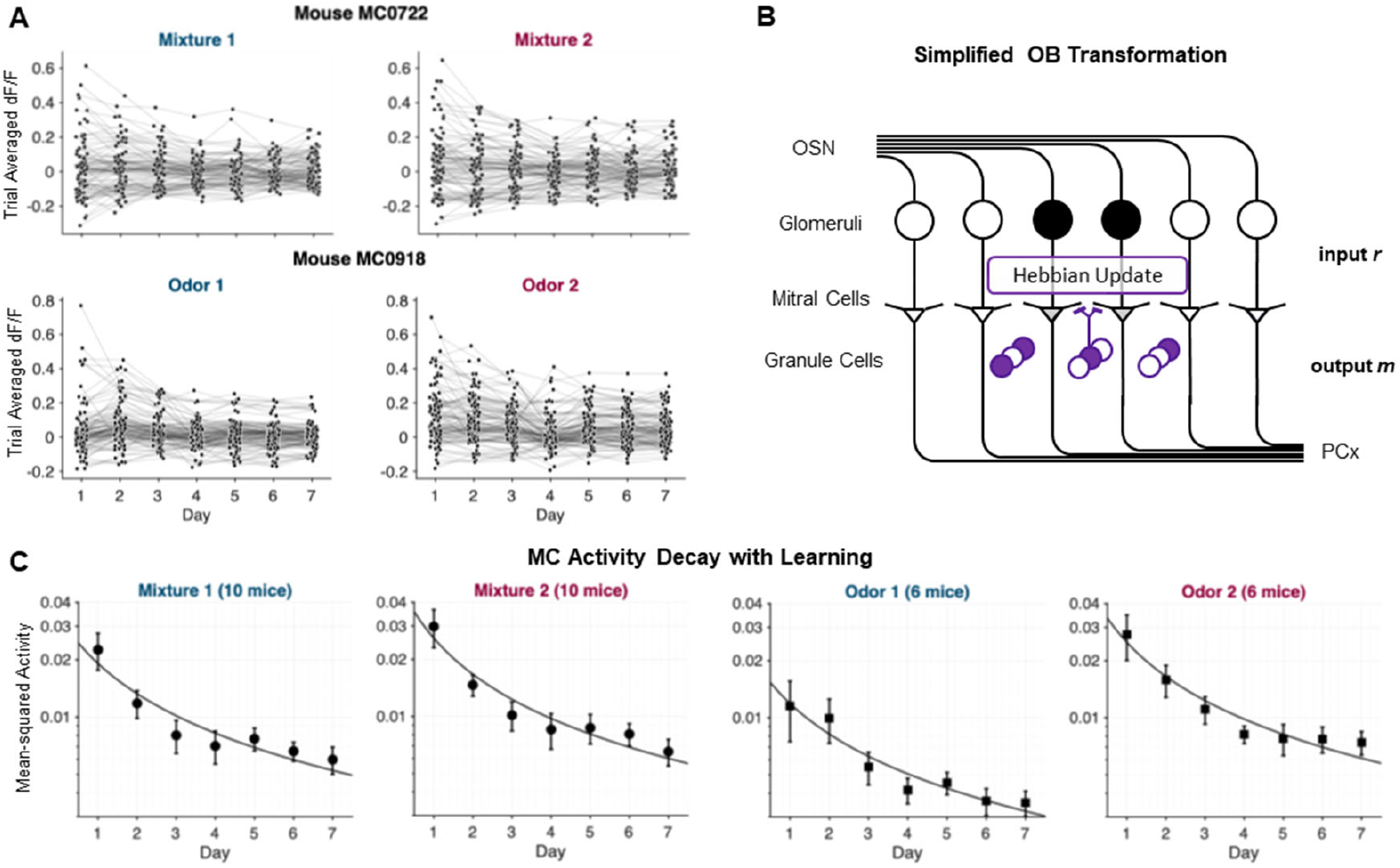
Gain adaptation in the MC population activity. **(A)** Single-neuron activity trajectories across days of odor exposure. Each dot represents the trial-averaged activity of an individual MC. Top panels show responses from a mouse exposed to a pair of similar odor mixtures (Mixture 1: 52% Heptanal/48% Ethyl Tiglate; Mixture 2: 48% Heptanal/52% Ethyl Tiglate). Bottom panels show responses from a mouse exposed to a pair of dissimilar odors (Odor 1: 100% Heptanal; Odor 2: 100% Ethyl Tiglate). **(B)** Simplified OB circuit model. MCs (triangles) receive excitatory input ***r*** from glomeruli (circles; filled = active, empty = inactive). Mitral cell population output ***m*** is sent to downstream areas including piriform cortex (PCx). GCs (purple circles, filled/empty -- active/inactive) form reciprocal dendrodendritic synapses with the MCs, providing feedback inhibition onto the MCs. **(C)** Comparison of model results with experimental data. In the mean-field model (solid curve), population activity decays as described by Equation 1. In the data (dark circles), mean-squared population activity (±SEM) across all mice for each odor condition follow similar dynamics.

To test this hypothesis quantitatively, we developed a computational mean-field model that predicts the temporal evolution of population activity (Supplementary Information Section 2). The model considers OSN activation represented by a vector ***r*** driving the population of MCs. The MCs are connected to a single GC that provides inhibitory feedback. This simplification is motivated by the sparseness of GC responses, which implies that very few GCs are responsive to any given odor. It also allows us to obtain a closed-form solution for the MC activity. The network implements the following mechanisms: (1) GC activity is proportional to weighted mitral cell responses, (2) MC outputs ***m*** are represented by the difference between OSN inputs and inhibitory feedback they receive from the GC, and (3) dendrodendritic synapses are modified via Hebbian plasticity. Since the observed population average of the MC activity is close to zero (Fig. S2A-D), in the model, we computed the expected variance of the MC population activity as a function of time measured in the number of odor presentations:

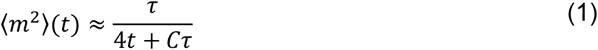

This equation predicts a power-law decay in population activity variance, where *τ* represents the characteristic timescale of synaptic plasticity and *C* is the fitting parameter that accounts for initial MC responses. We fit this model to the mean-squared MC activity of responsive neurons (on average 37 neurons per animal; Methods) across all recorded animals (Fig. 2C). We observed an agreement of the observed decay in the MC response variance with Equation (1). This agreement was observed across mice and odor conditions, with similar time constants *τ* recovered across experiments (Table S1; Fig. S2E-H for individual animal fits).

Overall, using experimental data of MC population responses to repeated odor presentations across several days, we observed a robust decrease in response amplitudes. This decrease was quantitatively described by the theoretical result [Equation (1)] derived using plasticity in dendrodendritic MC-GC synapses. Good agreement between experimental and theoretical results suggest that MC-GC plasticity may play a role in gain adaptation.

### Pattern separation and convergence in MC responses

The adaptation of MC population responses under long-term odor exposure differs between similar and dissimilar odor pairs. For highly similar odor mixtures presented over a week, the angle between MC population representation pairs progressively increased (Fig. 3C, light trace). At the same time, as shown in the previous section, the lengths of the population vectors decayed. As a result, the Euclidean distance between the representations of similar odor pairs by MCs remained approximately constant over time (Fig. 3A, left; Fig. S3A, light trace).

**Figure 3.**
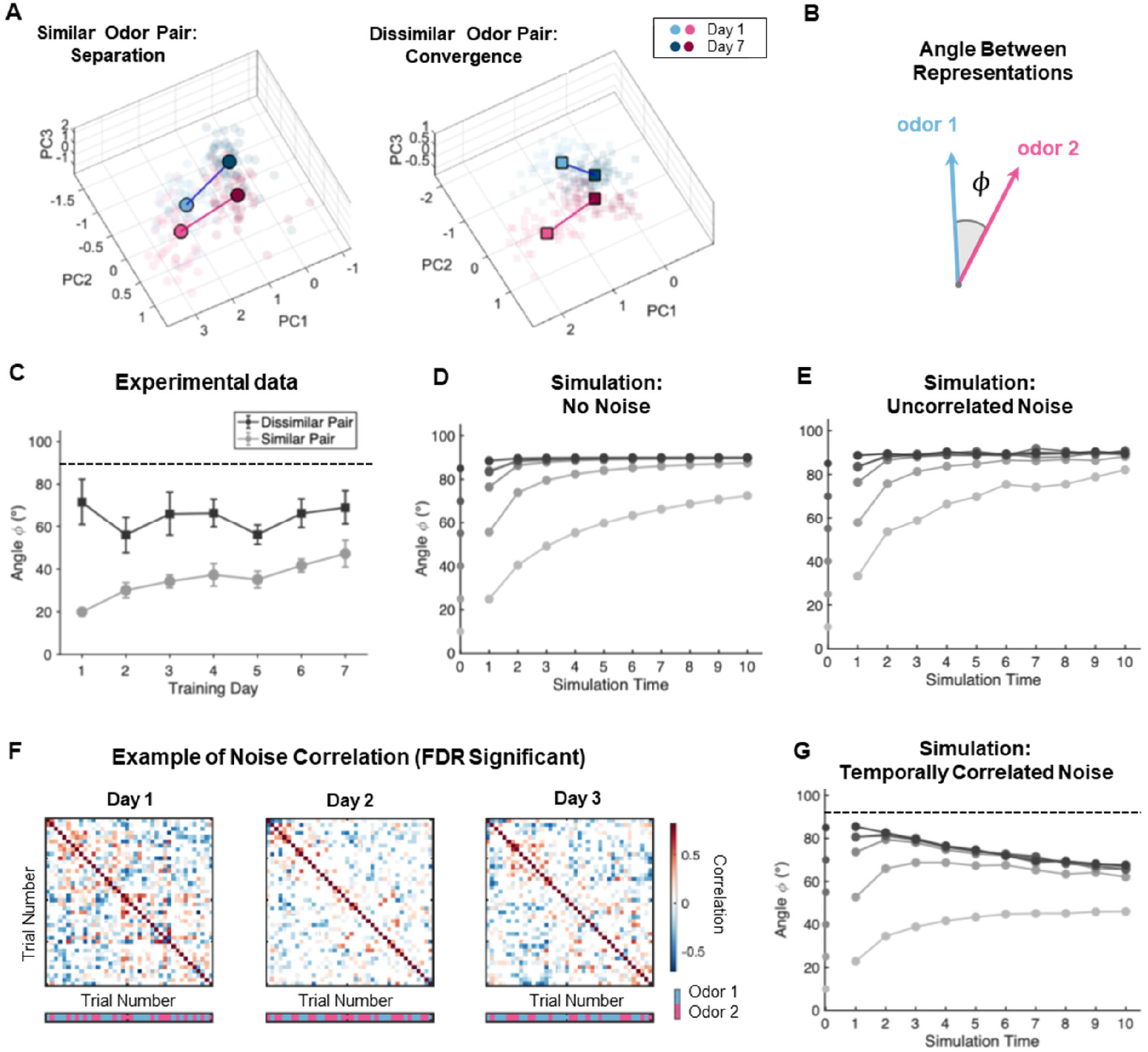
Pattern separation and convergence in OB odor responses. **(A)** Similarity-dependent MC activity changes in population responses. 3D visualization of PC1-PC3 of mitral cell population vectors across all trials from representative animals. Left: mouse exposed to similar odor mixtures (Mixture 1 vs. Mixture 2); Right: mouse exposed to dissimilar odors (Odor 1 vs. Odor 2). Transparent points show individual trials; solid points show daily trial averages. Data from Day 1 (light) and Day 7 (dark) illustrate opposing dynamics: similar odor representations separate over time while dissimilar odor representations converge. **(B)** Angle between odor representation vectors of two odors *ϕ*. **(C)** Mean angular separation (±SEM) between odor population vectors across training days. Similar odor pairs (gray) progressively separate, approaching a larger angle, while dissimilar pairs (black) remain at a non-orthogonal angle. **(D)** Model simulation with no noise. Evolution of the angle between odor pairs for varying initial similarities (grayscale levels represent different starting angles). Without noise, all odor pairs separate toward orthogonality regardless of initial similarity. **(E)** Model simulation with uncorrelated Gaussian noise. Same format as panel E, but with independent Gaussian noise (10% of input variance). Uncorrelated noise does not prevent separation toward orthogonality and fails to reproduce experimental convergence of dissimilar pairs. **(F)** Trial-to-trial noise correlations in experimental data. Correlation matrices of noise residuals (trial responses minus average response) for the first 40 trials per day from one representative animal. First three days shown. For each day, significant trial-by-trial correlations are displayed (two-tailed t-test, FDR corrected using Benjamini-Hochberg procedure); non-significant correlation entries are filled with white. Block structure along the diagonal reveals temporally correlated noise in adjacent trials. **(G)** Model simulation with temporally correlated noise. Same format as panels E and F, but with temporally correlated Gaussian noise (10% of input variance, adjacent-trial correlations). Correlated noise enables convergence of initially dissimilar representations to a finite asymptotic angle (dashed line), matching experimental observations.

For dissimilar odor pairs, the angle between population vectors remained approximately constant, though not equal to 90 degrees (Fig. 3C, dark trace). Here too, the lengths of the population vectors decreased, leading to an overall convergence of dissimilar odor representations (Fig. 3A, right; Fig. S3A, dark trace).

The functional consequences of such an adaptation depend on the readout mechanism implemented in MC target regions. For similar odor pairs, because the angle between representations diverges, normalization can make the downstream signals more distinct. For example, if downstream processing normalizes population vectors to equal lengths, adaptation causes representations of similar odors to move further apart. For dissimilar odor pairs, under this normalization scenario, the distance between representations remains unchanged. If, on the other hand, downstream regions rely on non-normalized population responses, we expect no divergence for similar pairs and a convergence for dissimilar pairs. Thus, the functional interpretation of these transformations depends on the nature of the readout mechanism.

We next interpreted these results in light of our computational model. In the model, a pair of input vectors ***r*** was delivered intermittently to a population of MCs, which interacted with a population of GCs via dendrodendritic synapses (Fig. 2B). The connection weights between MCs and GCs were modified according to Hebbian plasticity rule (Supplementary Information Section 1). The model then computed the output population vectors ***m*** resulting from the interdigitated exposure to odor pairs.

In our simulations, we used 10 MCs and 750 GCs and verified that the conclusions are robust to the number of neurons, provided the population size is sufficiently large (Supplemental Information Section 4). MCs inputs from the odorant receptors (represented by the vector ***r***) for each smell remained unchanged over days, while the responses on MCs combined in the vector ***m*** were modified by GC inhibition. The network was presented with pairs of input vectors ***r***, ranging from highly similar to highly dissimilar. Similarity within a pair was defined by the angle between the input vectors ***r***, which varied from 10° (similar pairs) to 85° (dissimilar pairs, Fig. 3D). We then monitored how the angle between the output vectors ***m***, representing MC responses, evolved as a function of exposure duration.

We observed that, for *all* odor pairs, both similar and dissimilar, the angle between MC representations systematically increased with odor exposure time (Fig. 3D). In our computational model, this angle asymptotically approached 90°, implying that odor representations became orthogonalized. This behavior was evident both when MC inputs from receptor neurons contained no noise (Fig. 3D) and when MCs, in addition to odorant-dependent inputs, received uncorrelated white noise (Fig. 3E; Methods; Table S2). Thus, our model explained the divergence of similar odor representations over time (Fig. 3C). This occurred because GC feedback learned the overlaps between odor pairs and subtracted them from MC responses, leading to orthogonalization (Supplemental Information Section 4.1). By contrast, in the experimental data, the angle between representations of dissimilar pairs never reached 90° and instead remained between 50° and 80° (Fig. 3C). This result was *inconsistent* with the predictions of our simple computational model (Fig. 3D, E).

To refine our model, we examined the structure of trial-to-trial noise in MC responses. In the experimental data, we observed substantial noise correlations across trials (Fig. 3F). Incorporating temporally correlated noise into our model successfully reproduced the convergence of initially dissimilar representations to a stable intermediate angle between 0° and 90° (Fig. 3G; Supplementary Information Section 4.2; Fig. S3C-D). These results demonstrate that realistic noise correlations, in addition to the learning rule, are essential for explaining the representational dynamics observed in the OB.

Overall, we showed that the angle between representations of similar odors by the MCs diverge over time. This behavior is replicated by the computational model which included Hebbian learning in the dendrodendritic MC-GC synapses. By incorporating a realistic temporal model for noise, the same model accounts for stable angle between representation of dissimilar odors. Together, these results point to a single circuit mechanism capable of generating two very distinct effects, depending on the relationship between the odor pairs.

### Structured connectivity may drive representational drift beyond pattern adaptation, separation and convergence

In the previous section, we described the relative transformations of representations of pairs of odors under repeated exposure. Here, we examine the global rearrangement of population vectors across several days. To quantify these global changes, we investigated the subspace spanned by the population response vectors to several odors and its evolution over time (Fig. 1B-D, shaded region). In the simplest form of odor adaptation, the two population vectors representing an odor pair move relative to one another within the same subspace (Fig. 1B, C). This type of adaptation simplifies the readout, since the same linear classifier can discriminate between the odors across days. Alternatively, the subspace itself may change, rotating as a result of odor exposure (Fig. 1D). Such a global transformation could complicate discrimination across days. We therefore analyzed how odor exposure affects the population subspace and its stability.

To characterize global transformations, we employed Jordan principal angles (JPAs) (Chae et al., 2019; Jordan, 1875), which quantify the geometric relationship between high-dimensional subspaces. Our goal was to compare the subspaces spanned by population vectors representing the same pair of odors across days. For two 2D planes in 3D space, a single JPA captures the geometric angle between these planes. In higher dimensions, the number of JPAs exceeds one; for instance, the relative position of two 2D planes in 4D space is described by two angles. Intuitively, the minimum (maximum) JPA corresponds to the smallest (largest) angle between any two vectors drawn from the two subspaces. Computing JPAs between odor representations across days provides a convenient way to measure population subspace rotations induced by adaptation.

We computed the minimum and maximum JPAs between population vectors representing responses to odor pairs on day 1 and later days for n = 16 mice. For both similar (Fig. 4B, n = 10) and dissimilar (Fig. 4C, n = 6) odor pairs, JPAs were consistently greater than zero, indicating subspace rotation. Notably, JPAs remain unchanged if population vectors rotate within the same plane. Thus, our findings cannot be explained solely by the relative transformations of odor representations described in the previous section. Instead, the presence of non-zero JPAs across days points to a global transformation of odor representations.

**Figure 4.**
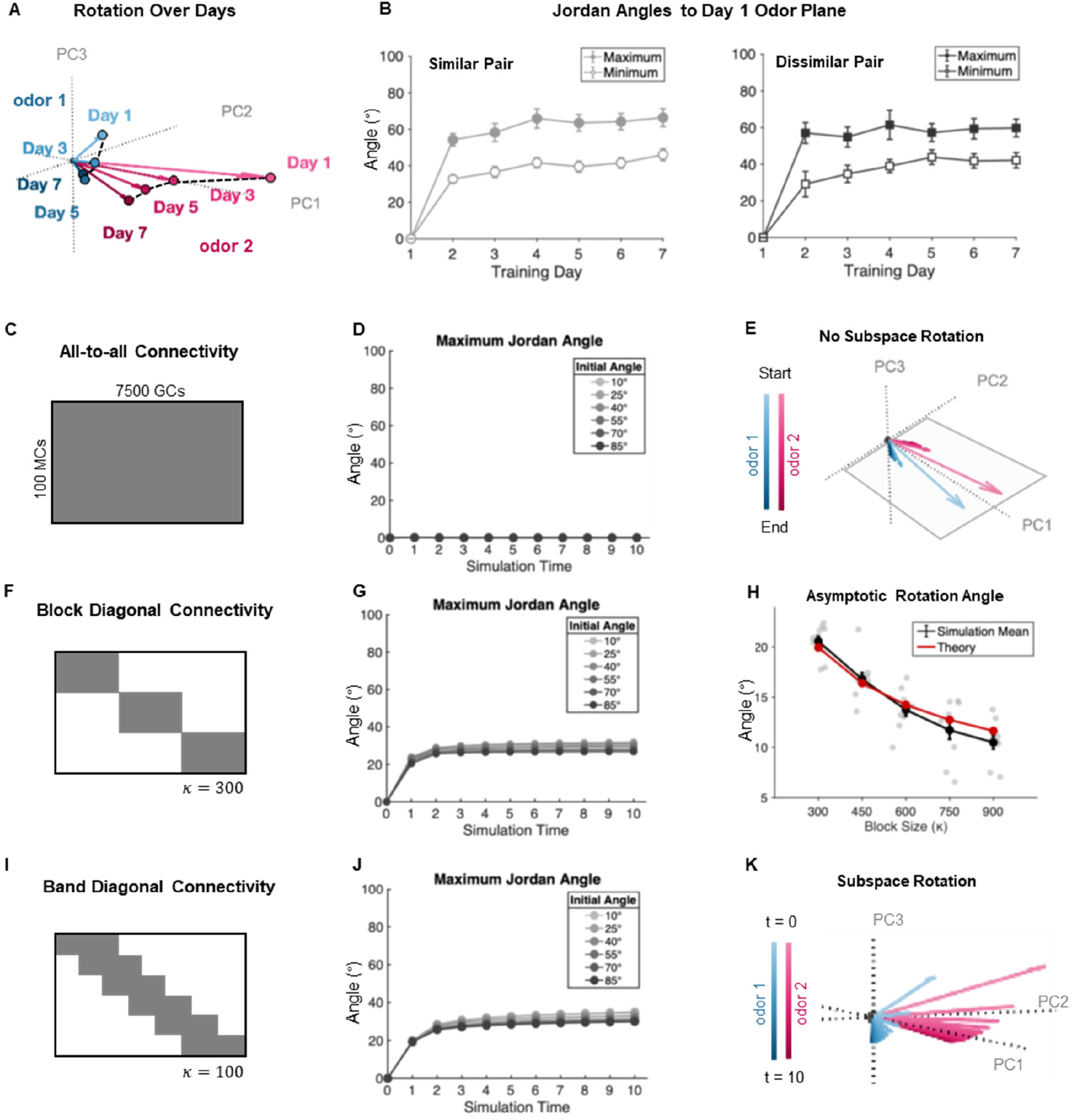
Representational drift results from structured connectivity in the MC-GC circuit. **(A)** Experimental evidence of representational drift. Principal component visualization of daily odor trial average from a representative mouse exposed to dissimilar odors. PCA performed on trial average of all days; data for every other day shown for clarity. Odor representations progressively occupy different subspaces over time, indicating systematic rotation distinct from decorrelation within the initial odor plane. **(B)** Quantification of subspace rotation using Jordan principal angles. Maximum and minimum Jordan angles between the Day 1 odor subspace and subsequent days measure how the two-dimensional odor representation plane rotates in neural space. Both angles increase progressively for similar (left) and dissimilar (right) odor pairs, indicating that the rotation magnitude is independent of odor similarity. Data show mean ± SEM across all animals. (**C, D, E**) Network simulations for fully connected MC-GC network. (C) MC-GC connectivity matrix (gray represents non-zero connections). (D) Jordan angles between planes defined by pairs of odor representations. For fully connected network, the representation plane does not rotate. (E) The dynamics of odor representations are visualized in PC space. No rotation is evident. (**F, G, H**) Network simulations for a sparse MC-GC connectivity. (F) In the block-diagonal connectivity matrix, most weights are fixed at zero (white, 90% of weights). The blocks of weights along the diagonal (gray) are allowed to change. (G) The maximum Jordan angle between planes defined by pairs of representations and the initial plane as a function of time. Sparse connectivity leads to the representation plane rotation. After certain time of exposure, representation plane stops rotating and the maximum Jordan angle reaches a steady value. (H) The asymptotic value of rotation angle of a single input as a function of block size. Rotation is weaker for a bigger block size. Simulation results (black, average over 10 random trials) agree well with theory (red, derived in Supplementary Information Section 3.4). (**I, J, K**) Network simulations for band-diagonal sparse MC-GC connectivity. (I) The GC-MC connectivity matrix. Gray regions indicate allowed connections (~3% of all weights). (J) As in the case of block-diagonal connectivity, the representation planes rotate after repeated odor exposure. (K) The dynamics of odor representations visualized in the PC space. The rotation of the representation plane is similar to data in (A).

This subspace change resembles representational drift reported in sensory cortices, where stimulus encodings change systematically over timescales of days to weeks (Bauer et al., 2024; Deitch et al., 2021; Marks & Goard, 2021). Piriform cortex data indicates that representational drift depends on odor presentation frequency: less frequently presented odors drift more. To evaluate the frequency dependence of representational drift in the OB, we examined the rotation of odor representations obtained with no or minimal odor exposure, estimated using a different previously published dataset (Shani-Narkiss et al., 2023; Shani-Narkiss et al., 2020). We found that, in the absence of odor exposure, population vectors drift to the extent that is not different with the case of daily odor exposure presented above (Fig. S4A). This observation suggests that OB drift does not depend on odor exposure frequency. This finding further suggests that the exposure frequency dependence observed in cortex may have cortical origins: because cortical and OB drift appear to have different dependences on odor presentation frequencies, the mechanism of the OB subspace rotation described above is likely not to be inherited from the cortex, despite substantial feedback projections from the cortex to the OB.

To understand the subspace rotation described above, we simulated the adaptation of MC responses to pairs of odorants using our model. We found that, in the case of all-to-all MC-GC connectivity, MC representation subspaces do not rotate (Fig. 4C-E). The lack of representation subspace rotation is observed for all dissimilar and similar odor pairs, suggesting that decorrelation preserves the original odor subspace (Fig. 4D). This finding holds true even in the presence of synaptic update stochasticity, which has been shown to drive drift in sensory and hippocampal systems (Fig. S4B; Supplementary Information Section 3.2) (Bauer et al., 2024; Eppler et al., 2025; Morales et al., 2025; Qin et al., 2023). Overall, our computational model suggests that with all-to-all structureless connectivity between MCs and GCs, Hebbian updates preserve the original odor subspace.

Next, we tested if the presence of structure in the MC-GC connectivity might explain the subspace rotation. Indeed, the OB spatial structure restricts GC dendrites to local interactions with nearby MCs, creating connectivity constraints that deviate substantially from all-to-all connectivity (Arnson & Strowbridge, 2017; Egger & Urban, 2006; Kim et al., 2011; Nagayama et al., 2014). We therefore tested whether structured connectivity patterns alone could reproduce the observed subspace rotation.

We first simulated the circuit model with the simplest form of structure, the block-diagonal connectivity (Fig. 4F). With such a structure, the model successfully generated subspace rotation, with the averaged over days Jordan angles comparable to experimental observations (maximum angle in Fig. 4G; minimum angles see Fig. S4C). The block-diagonal assumption also enabled analytical tracking of MC output vector dynamics, revealing that rotation depends on both input statistics and block size: smaller blocks and greater variability across odorant receptor inputs produced larger drift angles (Fig. 4H; Supplementary Information Section 3.4). Our analysis also shows how structure in MC-GC connectivity induces subspace drift. In the block-diagonal model, plasticity operates independently within each block, producing differential suppression of MC responses across blocks. This block-specific scaling modulates the MC population response, causing representations to rotate away from the original input direction. Overall, the block-diagonal connectivity model yields interpretable subspace rotation in MC population responses, consistent with experimental data.

We then tested a more realistic band-diagonal connectivity, which allows each GC to influence every MC through multi-synaptic pathways. This structure captures the spatially dispersed connectivity patterns of the OB (Arnson & Strowbridge, 2017; Egger & Kuner, 2021; Hong & Wilson, 2015; Willhite et al., 2006). This architecture similarly produces subspace rotation (Fig. 4I, J) with trajectories resembling experimental data (Fig. 4K). Together, these results demonstrate that realistic anatomical constraints on plasticity, not just the learning rule itself, are essential for representational dynamics in the OB.

While our analysis of subspace rotation demonstrates global changes in odor representations across days, it does not directly establish whether these changes affect decoding. To address this, we trained linear classifiers (SVMs) on responses from each day and tested their performance both within and across days. For both similar and dissimilar odor pairs, cross-day classification accuracy declined as the interval between training and testing increased, indicating that subspace rotation leads to representational drift (Fig. 5A, B, left) (Masset et al., 2022; Rule et al., 2019; Schoonover et al., 2021). For example, for similar odor pairs, classifier performance decreased from 71% for same-day training and testing to 56% when training and testing were separated by six days (Fig. 5A). This decline is comparable in magnitude to drift observed in the piriform cortex [(Schoonover et al., 2021); Extended Data Fig. 10f].

**Figure 5.**
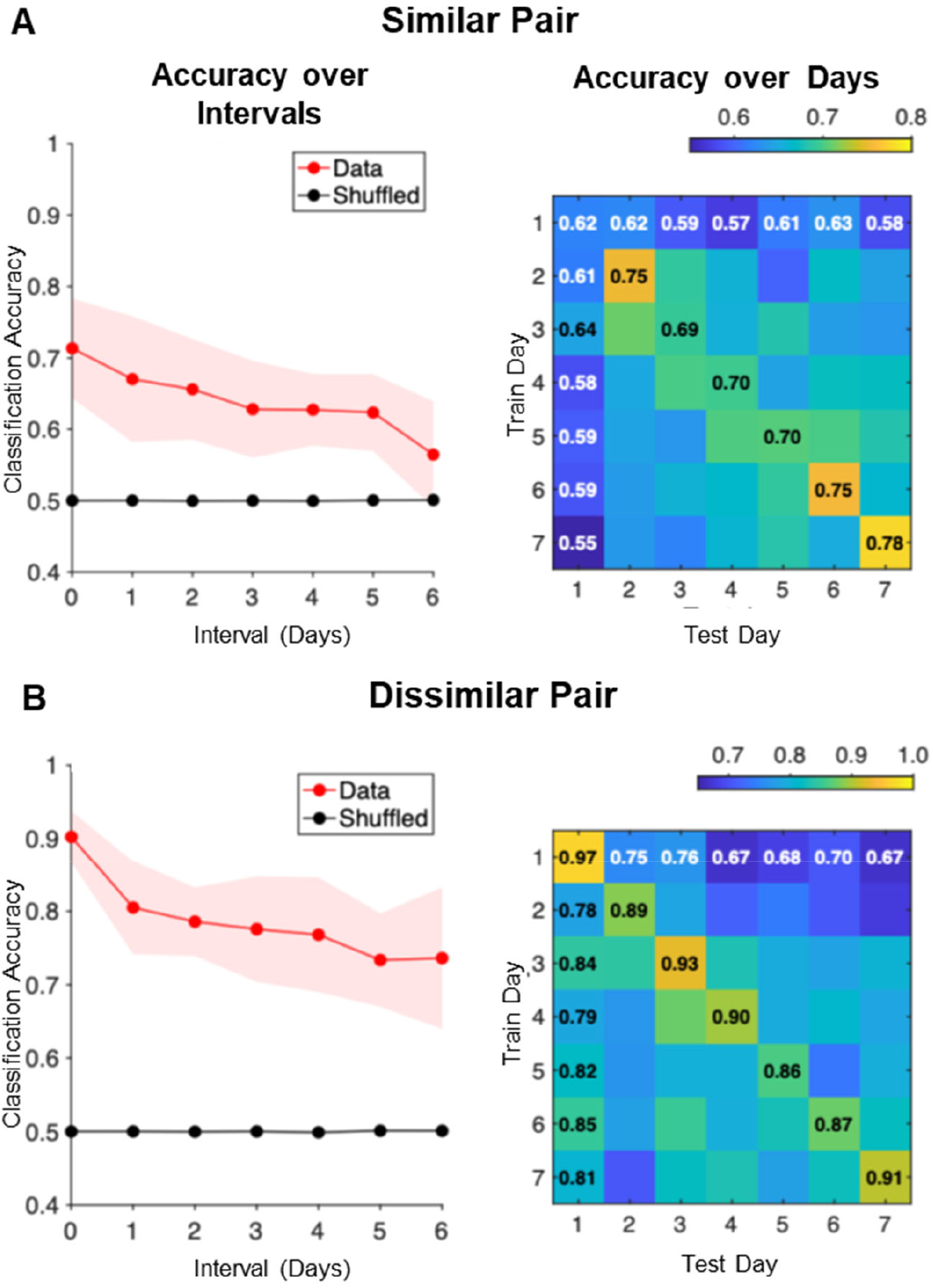
Classification accuracy (linear SVM) within and across days. (**A**) Classification accuracy for similar odor pairs. Within-day accuracy was obtained using leave-one-out cross-validation; across-day accuracy used models trained on all data from one day and tested on another day. Left: Classification accuracy as a function of time interval between train and test days. For interval equal to zero, within-day accuracy is shown. Mean ± SD across mice (n = 10). Right: Average classification accuracy matrix for all pairs of train and test days. Diagonal elements show within-day accuracy; off-diagonal elements show across-day accuracy. (**B**) Same as (A), but for dissimilar odor pairs (number of mice, n = 6).

At the same time, our classifier analysis revealed the impact of relative transformations between odor pairs described in the previous section. Within-day decoding of similar odor pairs improved at later days compared to Day 1, reflecting enhanced discriminability resulting from odor exposure (Fig. 5A, right, diagonal). This finding suggests that the pattern separation of similar odors increases their linear separability. By contrast, classifier performance for dissimilar odor pairs remained consistently high (Fig. 5B, right). Overall, these results indicate that global transformations of population vectors drive representational drift, necessitating adjustments to odorant classifiers downstream of the OB.

### Concentration gradient structure along representation manifolds is preserved during passive odor exposure

Our previous results suggest that representational change in the OB under repeated odor exposure can be decomposed into the following components: (i) population-level gain adaptation, (ii) relative pattern separation or convergence, and (iii) global subspace rotation. Although these components were derived from odor pairs, the same framework can be extended to larger stimulus sets. We therefore examined how *stimulus manifolds*, i.e. low-dimensional structures defined by MC population responses to mixtures of the same pure components, are encoded and transformed across days. In one scenario, the representations of individual mixtures are transformed independently across days, and their relative positions are scrambled. In this case, the angles between MC population vectors representing individual mixtures change, and the representation manifold undergoes a geometric deformation. We therefore refer to this form of representational change as *manifold deformation*. In another scenario, the representations of different mixtures rotate as a whole while preserving the relative angles between the MC population vectors. We refer to this hypothetical scenario as *rigid-body manifold rotation* (Pashakhanloo & Koulakov, 2023). Preserving representation manifolds may be important because it ensures that concentration gradients in stimulus space remain systematically reflected in population responses.

Testing the complete geometry of population vectors is not feasible, as it would require measuring odor responses from the entire ensemble of MCs. Indeed, even if population vectors undergo a rigid-body rotation during adaptation, observing only a fraction of the MC population would generally produce an apparent deformation. One possible exception arises when population responses form a one-dimensional (1D) manifold, such as that generated by responses to a binary mixture series. In this case, even partial sampling of the MC population may still reveal a robust rotation of the 1D manifold. We therefore examined 1D trajectories formed in by MC population responses to binary mixture series. We reasoned that if the ordering of mixture responses along the 1D trajectory, as determined by component concentration, is preserved across days, this observation would be consistent with rigid-body manifold rotation. Conversely, if mixture representations become scrambled along the 1D trajectory following adaptation, this would support the manifold deformation hypothesis. Thus, analysis of 1D binary mixture series may provide a first step toward inferring the geometric transformations of MC population representations induced by adaptation.

For a binary mixture of odors A and B, the component concentrations can be written as 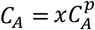 and 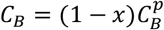, where 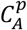 and 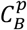 denote the concentrations of the pure components, and *x* parametrizes the mixture composition. The MC population response vector ***m*** depends on the component concentrations, ***m****=* ***m***(*C*_*A*_, *C*_*B*_), and thus becomes a function of the single parameter *x* for a given binary mixture series. As the relative concentrations of the components vary with *x*, the neural population vector ***m*** traces a 1D trajectory, in the population space, which we refer to as a *representation geodesic*. Under repeated exposure to the mixture series over multiple days, this representation geodesic is expected to change. Our goal is to examine whether the ordering of population vectors ***m*** corresponding to different mixture compositions along the geodesic is preserved across days. Previous studies have shown that the gradient between two pure odors projects orderly onto a one-dimensional axis in neural population space, supporting the existence of a representation geodesic (Adefuin et al., 2022; Carlsson et al., 2007; Fletcher, 2011). However, how such geodesics evolve under repeated exposure has not been investigated. We analyzed MC population responses from mice exposed to a binary mixture series of Heptanal and Ethyl Tiglate over more than a week (8 mixtures; see Table S1 for mixture composition). On Day 1, population responses were ordered according to mixture composition as visualized in the PC space (Fig. 6A). Across days, response amplitudes decreased, consistently with gain adaptation (Fig. S6A). The distribution of pairwise angles between odor population vectors became tighter over time, converging toward ~60° (Fig. S6B-C). Although the representation subspace rotated across days, the mixture ordering appears to remain generally preserved as assessed visually (Fig. 6A; Fig. S6D).

**Figure 6.**
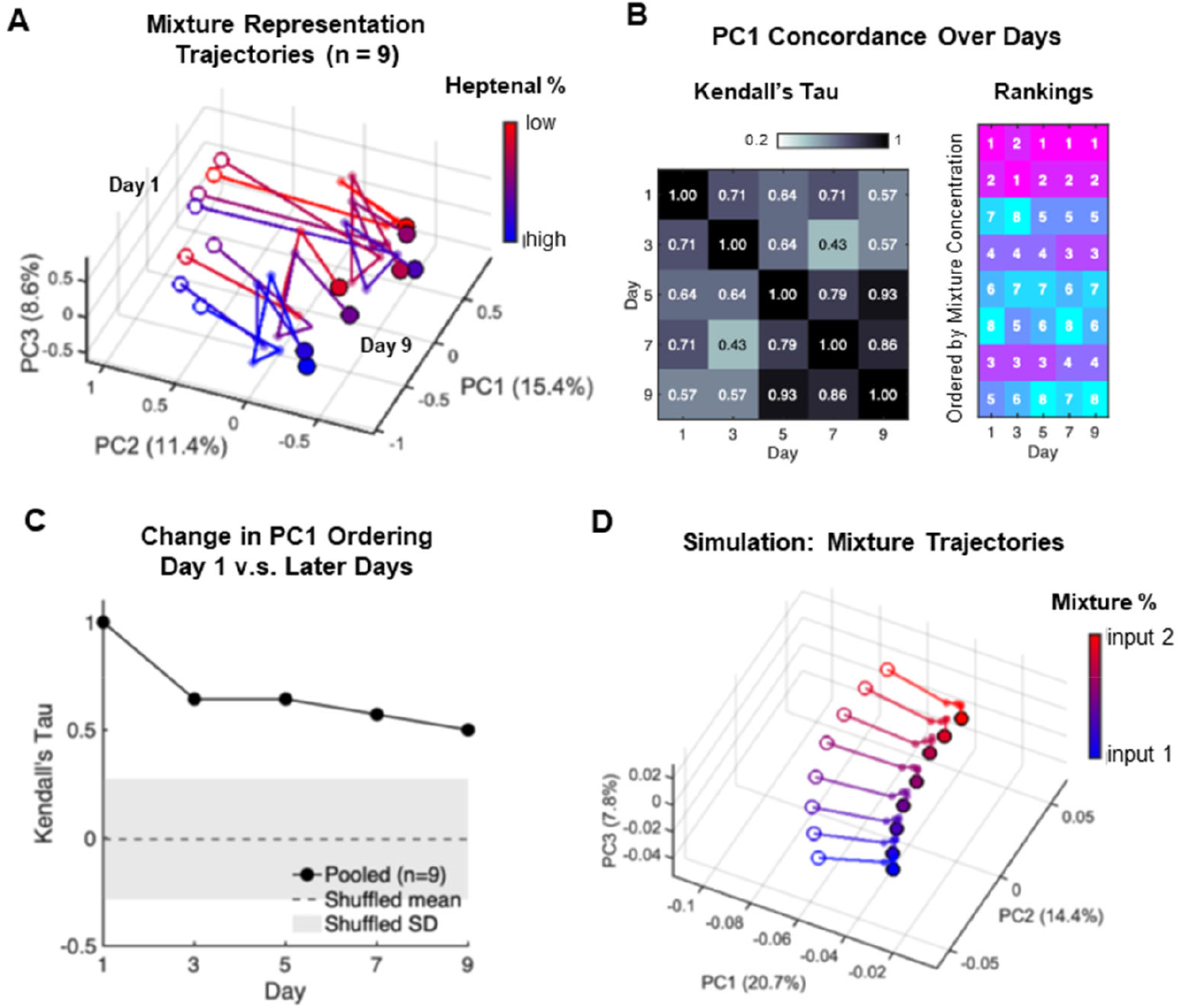
Representational drift preserves geometric structure of the 1D odor manifold. **(A)** Mixture representations maintain ordered trajectories during drift. Daily population vector (trial averaged) in response to 8 binary odor mixtures differing in the relative concentrations of Heptanal and Ethyl Tiglate in principal component space across all trials. Data collected on days 1, 3, 5, 7, and 9, with trajectories showing evolution from Day 1 (unfilled circles) to Day 9 (filled circles). Mixtures are color-coded by Heptanal content (red = low, blue = high) (Table S1). Despite substantial drift in PC space, the relative ordering along the concentration gradient remains consistent across days. **(B)** Concentration ordering is preserved in PC1 across days. Left: Kendall’s tau correlation coefficients measuring preservation of pairwise odor ordering between days. High correlations indicate maintained rank structure. Right: Rank ordering of the 8 mixtures along PC1 for each recording day, with colors representing rank positions (1 = highest PC1 loading, 8 = lowest). **(C)** Quantification of ordering preservation. Kendall’s tau comparing PC1 ordering on later days versus Day 1 shows significant conservation above chance levels (gray band shows shuffled control mean ± SD). **(D)** Model reproduces ordered drift dynamics. Simulated mixture trajectories using circuit model with block-diagonal connectivity and 10% input noise. Input mixtures created by linearly interpolating between dissimilar odor OSN population response vectors with proportions matching experimental angles on Day 1. Trajectories show trial-averaged evolution every 100 trials.

To quantitatively evaluate mixture ordering stability along the geodesic, we measured the rank stability of daily odor subspaces. To this end, we computed the first principal component (PC1) of the population response vectors for each day separately, which identifies the direction of the representation geodesic. This direction is expected to rotate across days due to the subspace rotation discussed above. We then projected mixture responses onto the corresponding PC1. If the ordering of representations across days is consistent or random, the ordering of mixtures along PC1 according to the mixture composition parameter *x* defined above is expected to be conserved or scrambled, respectively. Consequently, Kendall’s tau correlation of these orderings across days is expected to be close to one or zero, respectively. We found that Kendall’s tau correlations across days were consistently high (Fig. 6B, left), and the overall concordance across all days (Kendall’s W=0.87, p<10^-4^) indicated robust preservation of mixture ordering. Although occasional local inversions occurred between adjacent mixtures, the global rank order was significantly higher than in shuffled controls (Fig. 6C). Furthermore, the ordering axis aligned with the concentration gradient of the binary mixtures (Fig. 6B, right). Computational modeling incorporating structured connectivity and input noise reproduced these drift trajectories while preserving concentration ordering (Fig. 6D). Together, these results demonstrate that the low-dimensional structure of the stimulus ensemble (here, the binary mixture geodesic) is robustly preserved under representational change. Even as population responses undergo gain adaptation, pattern separation/convergence, and subspace rotation, the encoding of concentration gradients remains stable. This finding is broadly in line with a rigid-body-like rotation of the stimulus manifold, while the remaining local reordering of mixture representations and the tightening of the pairwise angle distribution across days (Fig. S6C) are indicative of additional non-rigid manifold transformations. Together, these results suggest that repeated odor exposure preserves the global organization of the representation manifold.

## Discussion

In this study, we examined how odor representations in the OB evolve over days of repeated passive exposure. By analyzing odor responses obtained using longitudinal calcium imaging in the bulb, we identified distinct but concurrent components of representational change: population-level gain adaptation, relative pattern separation or convergence, and global rotation of the encoding subspace. These phenomena could be collectively captured by a mechanistic computational model of the MC-GC circuit. The model demonstrates that dynamics and input-dependent plasticity within this recurrent network are sufficient to reproduce the spectrum of observed representational transformations.

Our first major finding concerns the overall activity of MCs during repeated exposure to similar and dissimilar odor pairs. Across days, we observed a gradual decay in the amplitude of population responses (Fig. 2A; Fig. S2A-D). Using a mean-field approximation of our circuit model, we showed that the variance of MC population activity on later days follows a power-law decay (Equation 1). This model accurately fits the population variance measured in our data across animals and odor conditions (Fig. 2C; Fig. S2E-H), indicating that activity-dependent plasticity in the MC-GC synapses can account for the long-term gain adaptation observed experimentally.

Second, we found that the relative changes between odor representations depend on the similarity of the odor inputs. For highly similar odor pairs, the angle between MC population vectors steadily increased across days, whereas for dissimilar pairs the angle remained stable around approximately a 70-degree angle (Fig. 3A, C). Using our model, we showed that Hebbian plasticity drives orthogonalization of input pairs regardless of their initial similarity (Fig. 3D, E). Incorporating temporally correlated noise, the presence of which we confirmed in the experimental data (Fig. 3F), stabilized the asymptotic angle between representations, producing convergence to a value below 90°, as observed experimentally (Fig. 3G).

Third, we observed a gradual rotation of the population encoding subspace across days, quantified using Jordan principal angles (JPAs). JPAs measured between the initial and subsequent days of odor exposure describes the degree of rotation of odor representation subspaces. Both similar and dissimilar odor pairs exhibited progressive increases in JPAs over time, indicating a global transformation of the representation plane (Fig. 4A, B). This subspace rotation also resulted in a decay of cross-day linear decoding accuracy (Fig. 5A, B). This implies that the linear decoder trained on day 1 of odor exposure could discriminate odors on subsequent days with lower accuracy. This finding suggests that subspace rotation in the OB may need to be compensated by a continual recalibration of downstream decoders, supporting its interpretation as representational drift. The subspace rotation could be accounted for by our MC-GC computational model through introducing biologically motivated structured connectivity constraints into the MC-GC circuit (Fig. 4F-H, J). Our model also predicted that repeated odor exposure leads to a plateau in rotation over time (Figs. 4G, J) as observed experimentally (Fig. 4B). By contrast, variability in the inputs alone, although capable of inducing some output rotation, did not generate the saturating rotation angle we observed experimentally (Fig. S5).

Finally, we extended the framework beyond the analysis of odor pairs to explain how odor representation manifolds defined by mixtures evolve under repeated exposure. One implication of our model is that plasticity and network dynamics in the MC-GC circuit should preserve the relative arrangement of the population vectors as absolute representations drift. This type of drift was termed here as rigid-body rotation (Pashakhanloo & Koulakov, 2023). In agreement with this hypothesis, the analysis of MC responses to an eight-odor binary mixture panel revealed that the ordering of representation vectors, when projected onto the first principal component, remained approximately stable across days (Fig. 6A-C). Simulations incorporating input noise and structured connectivity reproduced drift trajectories while maintaining the concentration ordering (Fig. 6D), demonstrating that the circuit can undergo substantial representational change without disrupting the low dimensional geometry of representation manifolds. Our findings are generally consistent with the global rigid-body rotation of the representation manifold, although some degree of local manifold deformation is also observed (Fig. S6).

While our model focuses on recurrent interactions between MCs and GCs, other structures in the OB could also contribute to the observed changes. Adaptation at the single-neuron level can occur through receptor desensitization (Antunes et al., 2014) or short-term synaptic plasticity during odor presentation or sniff bouts (Dietz & Murthy, 2005; Zhou et al., 2020). However, these mechanisms operate on short timescales (seconds to minutes) and are unlikely to account for the multi-day gain adaptation reported here. Additional gain control mechanisms are implemented in the glomerular layer by periglomerular cells and short-axon cells, which connect presynaptically to MCs or laterally among glomeruli. These interneurons have been reported to exert normalizing, population-wide gain control and their plasticity could, in principle, contribute to multi-day decreases in activity (Banerjee et al., 2015; Burton et al., 2017; Vaaga et al., 2017). However, our data show heterogeneous changes across individual MCs: some neurons decrease activity over days, while others increase (Fig. S1), a pattern more consistent with our model in which inhibitory strengthening is input and MC-specific. Nevertheless, we cannot rule out the possibility that the observed population-level changes arise from a combination of mechanisms across multiple interneuron types (Zavitz et al., 2020).

Numerous previous studies have modeled OB circuits (Cavarretta et al., 2018; Gilra & Bhalla, 2015; Kersen et al., 2022; Migliore et al., 2015; Migliore & Shepherd, 2008) to reveal how this network shapes inputs into the cortex. We propose that pattern separation in the OB over days arises from recurrent inhibition and plasticity at the MC-GC dendrodendritic synapses. In our model, local Hebbian plasticity progressively increases the angle between population vectors, orthogonalizing odor representations. Previous studies have shown that nonlinearity in neuronal responses can decorrelate input patterns via thresholding facilitated by sparse recurrent connectivity, providing a biologically plausible explanation for pattern decorrelation in the OB (Cayco-Gajic & Silver, 2019; Dasgupta et al., 2017; Wiechert et al., 2010). Several studies have proposed that learning and adaptation in the OB can further enhance pattern separation, especially for fine odor discrimination. For example, (Wick et al., 2010) compared feedforward and feedback (recurrent) inhibition and showed that the recurrent circuit produces stronger decorrelation. Linster and Cleland implemented a spike timing-dependent plasticity rule in a spiking model of vertebrate OB and insect antennal lobe networks, demonstrating that progressive sharpening of odor representations emerges from learning between principal neurons and their postsynaptic inhibitory partners, though they did not specifically examine the MC-GC circuit (Linster & Cleland, 2010). Several studies (Cecchi et al., 2001; Chow et al., 2012; Kersen et al., 2022) have built detailed anatomically and physiologically realistic models to study fine discrimination in the context of OB neurogenesis alone. These models show that OB neurogenesis helps decorrelate MC responses. Most recently, Meng and Riecke (Meng & Riecke, 2022) modeled plasticity between GCs and MCs during repeated exposure and assumed a non-monotonic relationship between synaptic spine formation rate and neuronal activity. In the context of input decorrelation, our work is consistent with these findings but differs in approach and further extends their results. By using a tractable linear system, we provide an analytical account of how population vectors evolve over the course of several days. We also propose that specific correlations in the MC input noise limit the levels of decorrelation in a manner consistent with experimental data.

The OB contains the first neural network that processes olfactory inputs and is generally thought to maintain relatively stable stimulus representations (Hirata, 2024), making our observation of representational drift at this early stage particularly noteworthy. Representational drift, on the other hand, has been previously reported in the piriform cortex (Morales et al., 2025; Schoonover et al., 2021). We found that, in the bulb, the degradation of linear classifier performance across days was numerically comparable to that reported in the cortex (Fig. S4A). Although we attribute the observed drift of the representation manifold to mechanisms within the OB, an alternative possibility is that the drift originates in the piriform cortex (Morales et al., 2025; Schoonover et al., 2021) and is conveyed to the bulb via corticobulbar feedback. Several arguments suggest, however, that the drift we observe may arise from intrinsic OB mechanisms. First, prior work indicates that during passive odor exposure, i.e. under the experimental conditions used here, cortical feedback remains relatively weak (Wu et al., 2020), making a cortical origin of OB drift unlikely. Second, both our experimental data and theoretical model show that the rotation of the MC representation plane quantifying the drift in the OB saturates after seven days of exposure, whereas cortical drift is reported to progress continuously without saturation. It is possible that longer-term structural connectivity changes in the OB driven by ongoing neurogenesis could in turn drive longer-term representational drift beyond the seven-day period, a possibility that warrants further investigation. Third, we find that the rate of drift in the OB is similar in overall magnitude to that seen in the cortex, but, unlike cortical drift, is not affected by odor exposure frequency. Overall, we propose that OB drift may be partially influenced by cortical feedback, but such feedback alone is unlikely to fully account for the phenomenon that we report.

At first glance, this aspect of drift appears to challenge classic neural coding theory, which emphasizes the stability of sensory representations in early processing stages (Morales et al., 2025; Patterson et al., 2013; Shani-Narkiss et al., 2023). To examine representation stability, we analyzed responses to a series of binary mixtures with gradually varying concentrations interpolating along a one-dimensional geodesic. We found that this geodesic rotated as a whole while preserving the relative positions of the population vectors representing individual mixtures. Preserving such a manifold may be important because it ensures that concentration gradients in stimulus space remain systematically mapped to population responses, even as plasticity alters their absolute configuration. Our computational model reproduced this coexistence of drift and manifold preservation, providing a mechanistic explanation for how stability and plasticity can co-occur in the OB.

Since representational drift was first described, most studies have emphasized stochasticity in synaptic weight updates as its primary source (Bauer et al., 2024; Eppler et al., 2025; Pashakhanloo & Koulakov, 2023; Qin et al., 2023). Here, we highlight a complementary mechanism, drift emerging from fixed structural constraints. In the OB, connectivity between MCs and GCs is both sparse and topologically constrained, and our model shows that such structure alone can drive systematic subspace rotations in the absence of noise (Fig. 4F-K). This rotation occurs from imperfect suppression of shared components between two odor representations due to the constraints on network connectivity. In a fully connected unconstrained MC-GC network, inhibitory feedback can selectively suppress overlapping components of activity patterns, and no subspace rotation occurs. By contrast, when each GC samples only a restricted subset of MCs, inhibitory feedback cannot selectively cancel shared components of two odor representations without also affecting their unique components. As a result, learning reduces overlap only approximately, leaving residual responses in the perpendicular to the original representation subspace directions. This residual activity manifests as a rotation of the encoding subspace, or representational drift (Fig. 4C-E). Similar ideas have been proposed elsewhere. For example, Kong et al. (Kong et al., 2024; Zabeh et al., 2025) suggested that differences in structural connectivity sparsity could account for the contrasting levels of drift observed in hippocampal CA1 versus CA3. The existence of structural constraints does not rule out stochastic fluctuations, but our findings suggest that, in the OB, fixed architecture may be a major contributor to drift. More broadly, such mechanisms may operate in other circuits where connectivity is locally organized. For instance, this may occur in primary visual cortex, where “like-to-like” connectivity is prominent. In this way, our results extend studies of representational drift by demonstrating that anatomical structure alone can generate long-term changes in population codes.

To further validate our proposed local learning mechanism between MCs and GCs, more studies of long-term synaptic plasticity at the dendrodendritic synapse are needed (Abraham et al., 2010; Egger & Kuner, 2021; Gao et al., 2009; Satou et al., 2005). Such plasticity would provide a compelling local mechanism for unsupervised perceptual learning. In sensory systems, decorrelation of neural representations for similar inputs can facilitate downstream classification by simple readout mechanisms, thereby accelerating subsequent task learning, as shown in visual cortex (Fink et al., 2025; Zhong et al., 2025). We hypothesize that analogous unsupervised changes in the OB may form the basis for reward association, odor identity reconstruction and other behaviorally relevant learning in the olfactory system (Hiratani & Latham, 2020; Krishnamurthy et al., 2022; Zavatone-Veth et al., 2023). These processes likely involve stronger and more specific cortical feedback than those studied here and may require considering the piriform cortex and OB together as an integrated circuit. Further work will be needed to integrate these mechanisms across bulb and cortex to understand how long-term learning shapes olfactory processing (Kumar et al., 2021; Wu et al., 2020; Yu et al., 2025).

Taken together, our findings provide a framework for understanding how the OB adapts to repeated odor exposure. By integrating quantitative analysis of longitudinal calcium imaging with a mechanistic MC-GC circuit model, we show that population-level gain adaptation, pattern separation, and subspace rotation can emerge from local plasticity and structural constraints. This work extends the concept of representational drift to an early sensory stage while demonstrating that low-dimensional manifold structure can be preserved despite ongoing change. More broadly, our results illustrate how stability and plasticity can coexist in recurrent sensory circuits and suggest that similar principles may govern other brain regions with sparse connectivity.

## Acknowledgements

This work was supported by the National Institutes of Health BRAIN Initiative grant number U19NS112953 and, in part, by National Science Foundation grant PHY-1748958 to the Kavli Institute for Theoretical Physics. C.X.Z. was supported by William R. Miller Fellowship at the School of Biological Sciences at Cold Spring Harbor Laboratory. S.N. thanks funding from the Simons Center for Quantitative Biology at Cold Spring Harbor Laboratory.

## Methods

### Experimental Dataset Details

#### Chu et al., 2016

Two-photon calcium imaging for mitral cell-specific calcium indicator was performed using longitudinal recordings of the same mitral cell populations over 7 consecutive days. During passive exposure sessions, mice experienced odorant presentations (4 s duration) of either a dissimilar pair (Odor 1, Heptanal; Odor 2, Ethyl Tiglate) or a similar pair ((mixture 1, 52% Heptanal/48% Ethyl Tiglate; mixture 2, 48% Heptanal/52% Ethyl Tiglate)) or followed by 15 s inter-trial intervals, with 75 trials per odorant delivered pseudo-randomly per session. No behavioral task or water reward was provided during passive exposure.

#### Li et al., 2018

Two-photon calcium imaging was performed every other day using longitudinal recordings of the same mitral cell populations across multiple sessions over 9 days. During passive exposure sessions, mice experienced odorant presentations (4 s duration) of eight different binary mixtures (54% Heptanal/46% Ethyl Tiglate through 46% Heptanal/54% Ethyl Tiglate in 2-3% increments), followed by 15 s inter-trial intervals, with 150 trials delivered pseudo-randomly per session. No behavioral task or water reward was provided during passive exposure.

#### Shani-Narkiss et al., 2020 & Shani-Narkiss et al., 2023

Two-photon calcium imaging was performed on awake head-fixed mice using longitudinal recordings of the same mitral cell populations across two sessions separated by 4 weeks. During odor presentation sessions, mice experienced odor panel presentations (2 s duration), in which Ethyl Tiglate was included, presentation is followed by 26 s inter-stimulus intervals, and each session contains 5 trials per odor. No behavioral task or water reward was provided during passive exposure. This dataset served as a control condition examining long-term stability of olfactory responses without chronic odor exposure.

### Data Analysis

#### Calcium Activity and Population Vector Formation

Following Chu et al. (2016) and Li et al. (2018), for each trial and each mitral cell, the calcium signal time series was normalized to the average fluorescence value during the baseline period (5 s before odorant onset) to calculate dF/F. Mitral cells that are classified as responsive in at least one session are considered (See criteria in (Chu et al., 2016)). The MC population response in each trial was expressed as an activity vector, which was a concatenation of the time-averaged dF/F from the first and second 2 s windows during the 4 s odorant period of each mitral cell. We used the trial average within each day’s session as the representation of odor response on that day. Using the full 4 s period to form population vectors did not change the main conclusions.

For Shani-Narkiss et al. (2020, 2023) data, we followed the original paper’s method to calculate relative fluorescence change (dF/F), defining baseline fluorescence for each cell in each trial as its mean fluorescence measured 5-2 s before odor onset. All traces were smoothed with a moving average filter (span = 5). To form the population vector, each entry represents the average dF/F during the 2-second odor presentation.

#### Principal Component Analysis

For individual mice, principal component analysis was performed for response vectors of all trials across all recording days in the Chu et al. (2016) and Li et al. (2018) datasets. For analysis and visualization across mice, a matrix of trial response vectors was created using all mitral cells that are classified as responsive in at least one session for each mouse using the first n trials (determined by the session with the lowest number of trials for a single odorant). Pooled response vectors were created by concatenating response vectors across animals. Given that calcium responses represent magnitude changes from baseline (dF/F) and mitral cell activation patterns are naturally sparse, PCA was performed without centering to preserve the meaningful zero baseline and activation magnitudes.

#### Jordan Angle between Subspaces

To quantify changes in odor representation geometry across days, we computed Jordan angles between odor subspaces. For each recording day, we defined the odor subspace as that spanned by the trial-averaged population vectors for each odorant. Specifically, for day *i* with *k* odorants, the subspace *S*_*i*_ was spanned by *k* vectors, where each vector represents the mean population response to one odorant across all trials on that day. Jordan angles between subspaces *S*_*i*_ and *S*_*j*_ (corresponding to different recording days) were computed using the following procedure. Let *A* and *B* be matrices whose columns are orthonormal bases for subspaces *S*_*i*_ and *S*_*j*_, respectively. The Jordan angles *ϕ*_1_, *ϕ*_2_,…, *ϕ*_*k*_ are defined as:

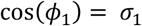

where *σ*_*1*_ are the singular values of *A*^*T*^ *B*, obtained from the singular value decomposition:

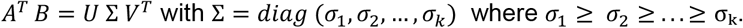

For the two-odorant datasets (Chu et al., 2016), this yielded two principal angles per day-pair comparison: a minimum angle (*ϕ*_*min*_ *= ϕ*_*1*_) and maximum angle (*ϕ*_*max*_ *= ϕ*_*2*_). For the eight-odorant dataset (Li et al., 2018), eight principal angles were obtained, representing the full spectrum of geometric relationships between the higher-dimensional odor subspaces.

#### Linear Classifier Analysis for Odor Pairs

Binary classification was performed using a linear Support Vector Machine with soft margins. Features were standardized (z-scored), and the soft margin parameter C was set to 1. For within-day classification, we used leave-one-out cross-validation, testing classifier prediction on each trial that was left out. Accuracy was reported as the average across all left-out trials. For across-day classification, we trained the SVM on all trials from one day and tested on all trials from another day. Performance on shuffled data was assessed by randomly permuting the odor labels (n = 1000 shuffles).

### Modeling Details

#### Model Fitting

We fit the mean-field decay model equation (1) to mitral cell activity in both individual animals and pooled populations across animals. For each responsive neuron, we calculated the squared trial-averaged activity and obtained the mean squared activity across the population for each day. The daily mean squared activity was then fit using least squares regression to estimate model parameters (Table S1).

#### Simulation

We initialized the network with random small weights and assumed fast dynamics for mitral cell-granule cell interactions coupled with slow Hebbian learning and weight decay (detailed modeling set-up see Supplementary Information Section 1). Inputs ***r*** are drawn from zero-centered Gaussian distribution.

The simulation protocol consisted of two phases:

- *Pre-exposure phase*: The network was presented with random inputs drawn from the same distribution as the experimental inputs to ensure proper weight initialization before chronic exposure
- *Chronic exposure phase*: Fixed input patterns were presented in an interleaved manner over many trials, mimicking the experimental paradigm

Network metrics, including principal angles between output representations, were evaluated every 100 trials for two-odor inputs and every 50 trials for eight-odor input panels, chosen to approximate the trail numbers in the experimental data. Input pattern (***r***), input noise, and weight update noise were drawn from Gaussian distributions. Detailed parameter values are provided in Table S2.

## Supplementary Tables

**Supplementary Table 1.**
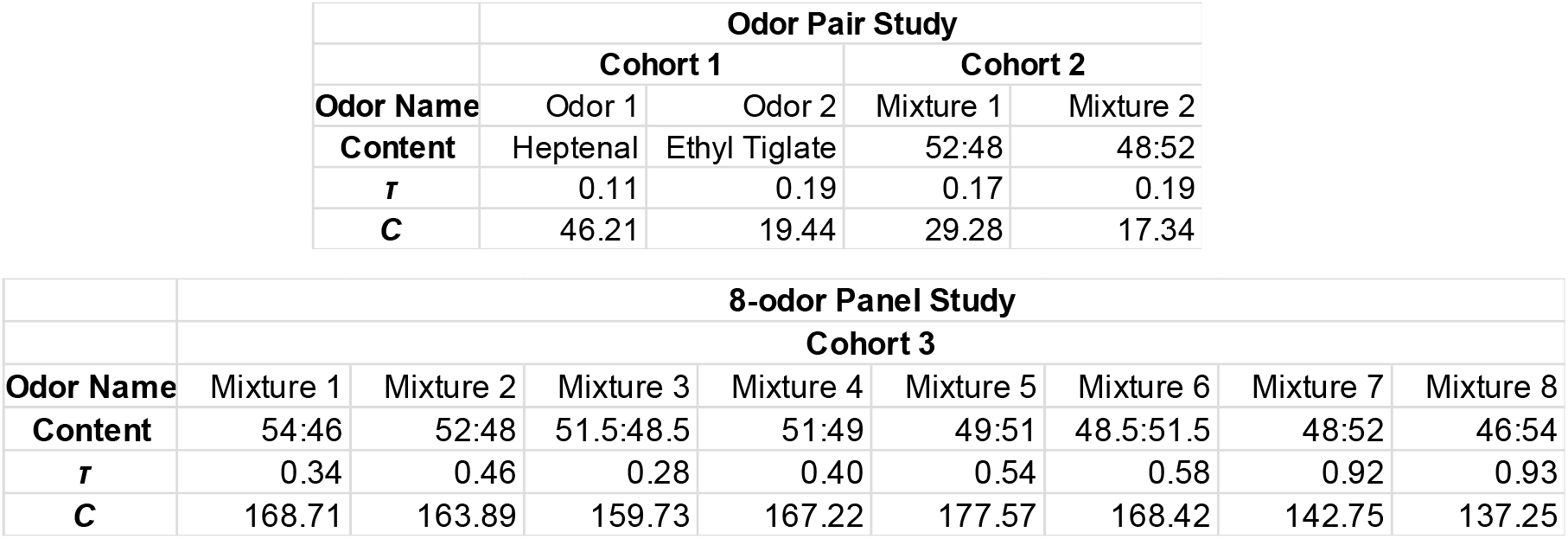
Mean filed decay fit. The average mean-field fitted parameters *τ* and *C* for each cohort exposed to each odor.

|  | Odor Pair Study |  |  |  |
| --- | --- | --- | --- | --- |
|  | Cohort 1 |  | Cohort 2 |  |
| Odor Name | Odor 1 | Odor 2 | Mixture 1 | Mixture 2 |
| Content | Heptenal | Ethyl Tiglate | 52:48 | 48:52 |
| $\tau$ | 0.11 | 0.19 | 0.17 | 0.19 |
| C | 46.21 | 19.44 | 29.28 | 17.34 |

Supplementary Table 1. Mean field decay fit.
|  | 8-odor Panel Study |  |  |  |  |  |  |  |
| --- | --- | --- | --- | --- | --- | --- | --- | --- |
|  | Cohort 3 |  |  |  |  |  |  |  |
| Odor Name | Mixture 1 | Mixture 2 | Mixture 3 | Mixture 4 | Mixture 5 | Mixture 6 | Mixture 7 | Mixture 8 |
| Content | 54:46 | 52:48 | 51.5:48.5 | 51:49 | 49:51 | 48.5:51.5 | 48:52 | 46:54 |
| $\tau$ | 0.34 | 0.46 | 0.28 | 0.40 | 0.54 | 0.58 | 0.92 | 0.93 |
| C | 168.71 | 163.89 | 159.73 | 167.22 | 177.57 | 168.42 | 142.75 | 137.25 |

**Supplementary Table 2.** Circuit simulation parameters. ***M***: Number of mitral cells (MCs) in the network. ***N***: Number of granule cells (GCs) in the network. ***θ***: Linear gain parameter relating GC membrane voltage to firing activity. ***η***: Hebbian learning rate controlling the magnitude of synaptic updates. ***λ*:** Synaptic decay (turnover) rate enforcing weight regularization. ***ε***_***ind***_: Independent input noise magnitude; Gaussian white noise applied independently to each neuron, with variance proportional to input strength. ***ε***_***sh***_: Shared input noise magnitude; Gaussian white noise shared between inputs, with variance proportional to input strength. ***σ***: Synaptic noise amplitude in Hebbian updates; Gaussian white noise with standard deviation calibrated to be on the same order of magnitude as the average absolute deterministic Hebbian update. ***κ*:** in block constraint, GC block size (columns per block); MC block size set proportionally. In band constraint, represent half-width (in GC indices) around each row’s center, with periodic wrap-around.

| | $M$ | $N$ | $\theta$ | $\eta$ | $\lambda$ | $\varepsilon_{ind}$ | $\varepsilon_{sh}$ | $\sigma$ | $\kappa$ |
| --- | --- | --- | --- | --- | --- | --- | --- | --- | --- |
| Fig. 3D | 10 | 750 | 0.5 | 0.05 | 0.0001 | 0 | 0 | 0 | / |
| Fig. 3E | 10 | 750 | 0.5 | 0.05 | 0.0001 | 0.1 | 0 | 0 | / |
| Fig. 3G | 10 | 750 | 0.5 | 0.05 | 0.0001 | 0 | 0.1 | 0 | / |
| Fig. 4C | 40 | 3000 | 0.5 | 0.05 | 0.001 | 0 | 0 | 0 | / |
| Fig. 4F | 40 | 3000 | 0.5 | 0.05 | 0.001 | 0 | 0 | 0 | 300 |
| Fig. 4I | 40 | 3000 | 0.5 | 0.05 | 0.001 | 0 | 0 | 0 | 100 |
| Fig. 6 | 20 | 1500 | 0.5 | 0.1 | 0.012 | 0 | 0 | 0 | 300 |
| Supp. Fig. 3C | 10 | 750 | 0.5 | 0.05 | 0.0001 | 0.05 | 0 | 0 | / |
| Supp. Fig. 3D | 10 | 750 | 0.5 | 0.05 | 0.0001 | 0 | 0.05 | 0 | / |
| Supp. Fig. 4C | 10 | 750 | 0.5 | 0.05 | 0.0001 | 0 | 0 | 0.01 | / |
| Supp. Fig. 5A | 40 | 3000 | 0.5 | 0.05 | 0.0045 | 0 | 0 | 0 | 300 |
| Supp. Fig. 5C | 40 | 3000 | 0.5 | 0.05 | 0.0045 | 0.1 | 0 | 0 | / |
| Supp. Fig. 5E | 40 | 3000 | 0.5 | 0.05 | 0.0045 | 0.1 | 0 | 0 | 300 |

## Supplementary Figures

**Supplementary Figure 1.**
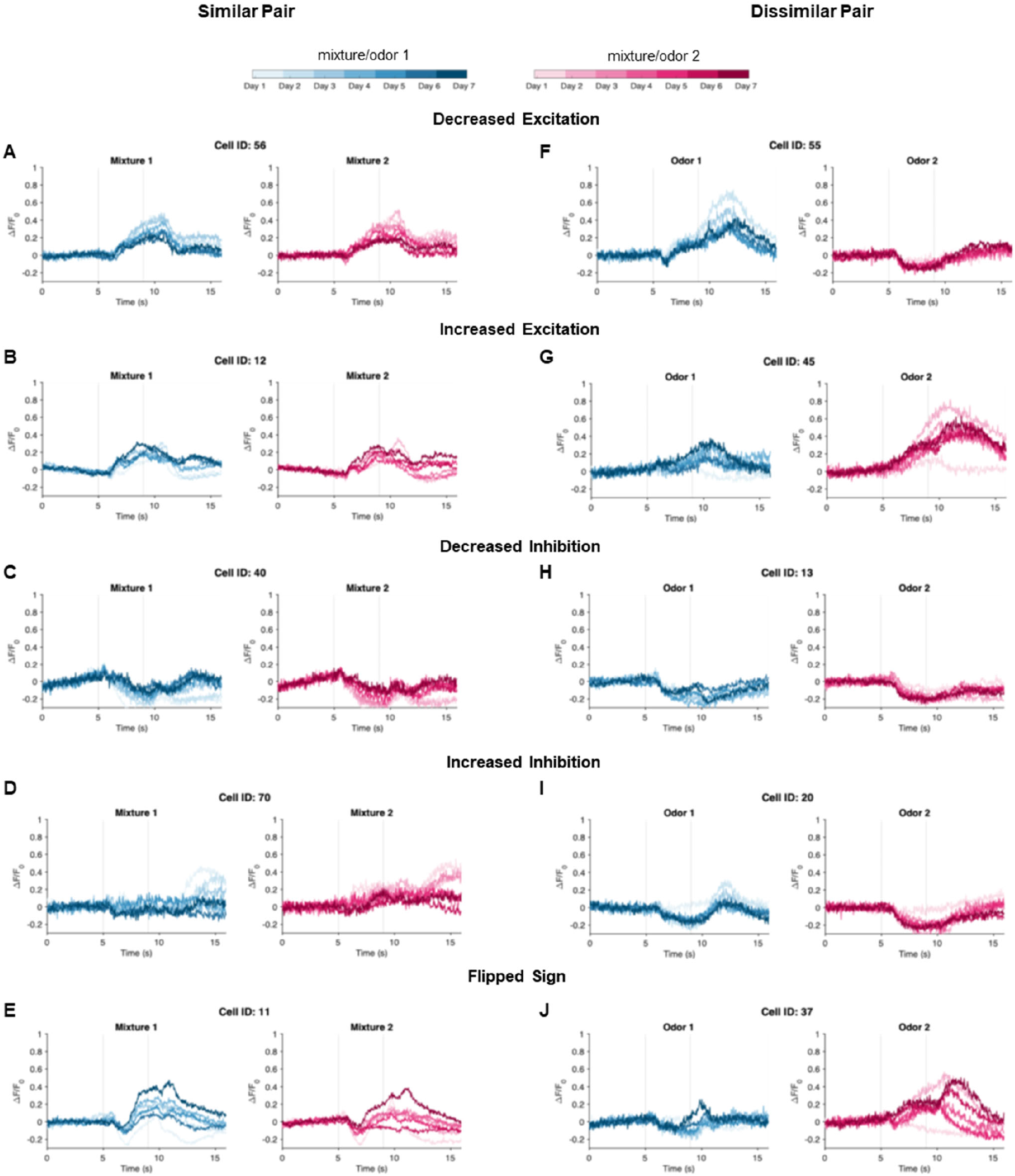
Individual mitral cells show diverse response dynamics during learning, despite population-level gain adaption. Trial-averaged dF/F traces are shown for individual neurons across seven days of training. Dotted lines indicate the 4-second odor presentation period. (**A-E**) Similar odorant pair during passive exposure. Representative examples of different plasticity patterns observed in individual mitral cells from one mouse. Neurons were selected to illustrate the range of response changes for Odor 1 during the odorant delivery period. Note that responses to Odor 2 may follow different temporal dynamics than Odor 1 responses in the same cell. (A) Decreased excitation: progressive reduction in excitatory responses over training days. (B) Increased excitation: enhanced excitatory responses over training days. (C) Decreased inhibition: reduction in inhibitory responses over training days. (D) Increased inhibition: enhanced inhibitory responses over training days. (E) Sign reversal: neuron switching from inhibitory to excitatory responses over training days. (**F-J**) Dissimilar odorant pair. Representative examples showing diverse activity pattern changes as in panels (A-E). These examples demonstrate that while the population shows overall decreased activity (Fig. 2), individual neurons exhibit heterogeneous response dynamics that cannot be predicted solely from population-level changes.

**Supplementary Figure 2.**
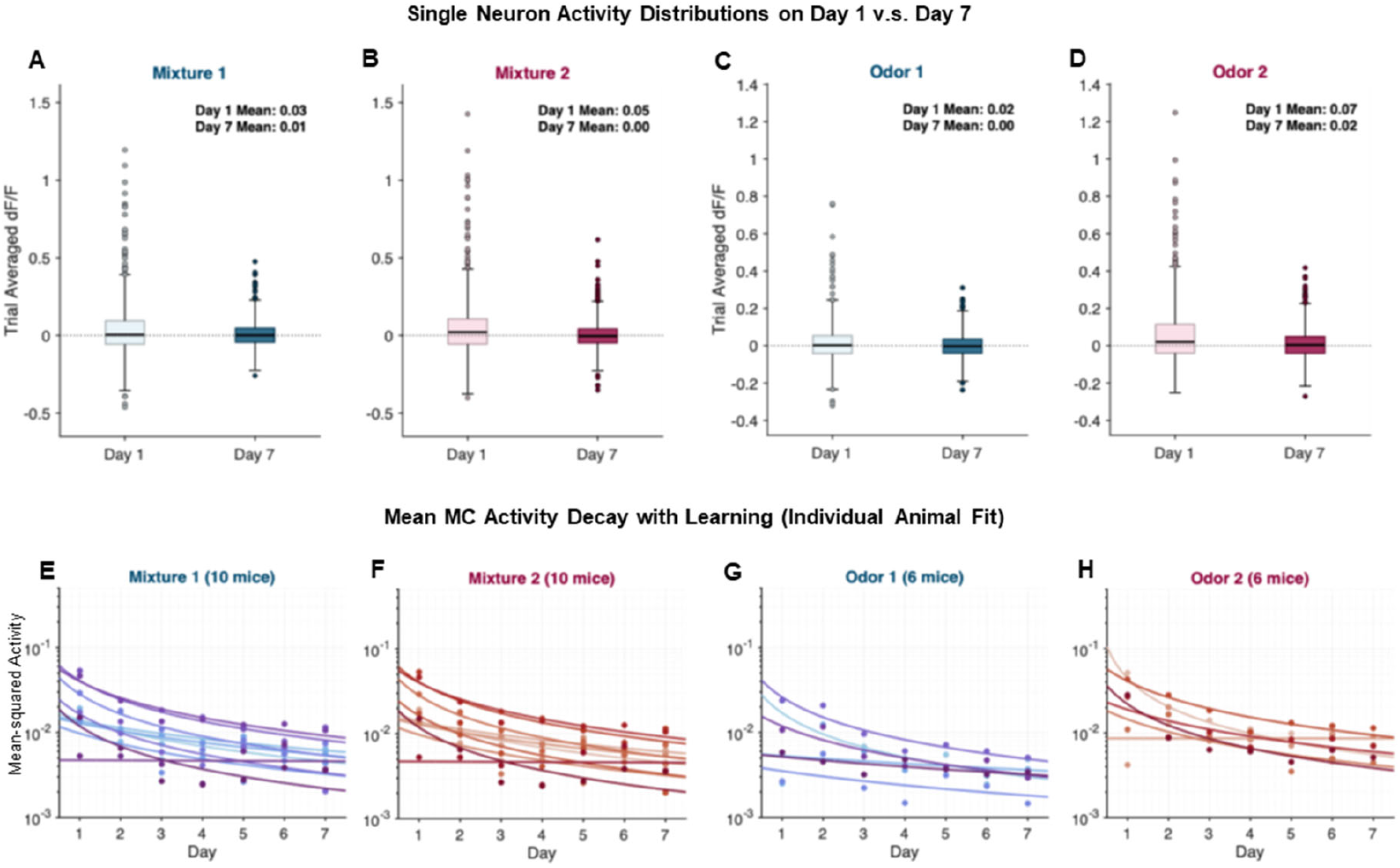
Population activity decay manifests as reduced dynamics on a population level, and the trend is consistent over mice. (**A-D**) Single-neuron activity distributions on Day 1 versus Day 7. Trial-averaged dF/F responses for individual neurons pooled across all animals exposed to each odor condition. Box plots show median, 25th and 75th percentiles; whiskers extend to 2× interquartile range. Note that mean activity remains near zero for all odorants, with neurons showing both excitation and inhibition relative to baseline, but the overall dynamic range decreases substantially by Day 7. (**E-H)** Individual animal fits to mean-field decay model. Each color represents data from one animal. Points show daily mean-squared population activity; lines show best-fit power-law decay functions.

**Supplementary Figure 3.**
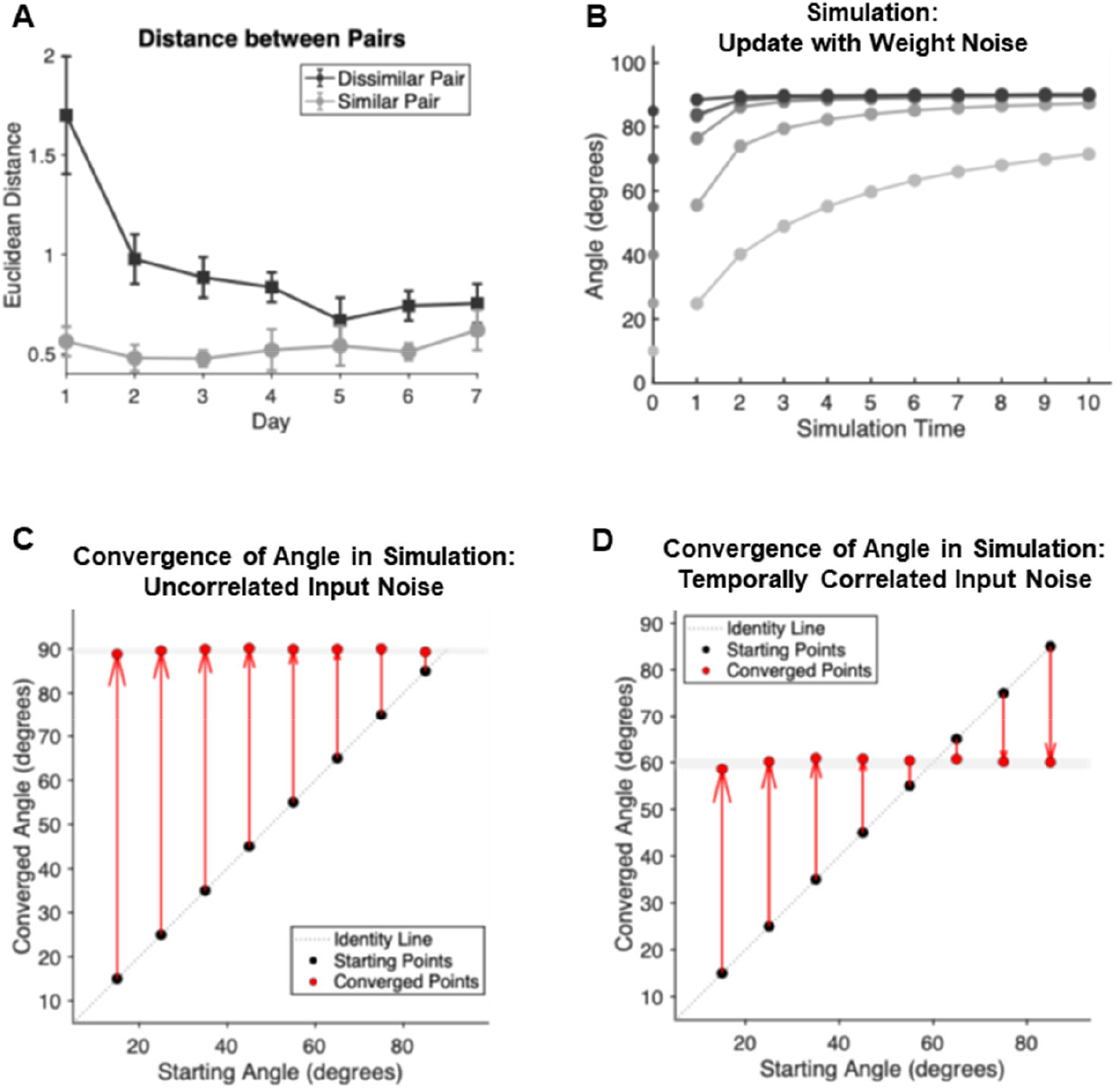
Angular dynamics depend on input similarity and noise correlation structure. **(A)** Between-odor Euclidean distance changes reveal similarity-dependent dynamics. During passive exposure, Euclidean distance between population vectors for dissimilar odor pairs (dark) decreases substantially over days, while distance between similar odor pairs (light) remains relatively stable despite overall activity decay (see Fig. 2, Fig. S2). Data show mean ± SEM across animals. **(B)** Weight update noise does not affect decorrelation dynamics. Simulation with random normal noise added to Hebbian weight updates (magnitude comparable) shows that decorrelation toward complete orthogonality remains unchanged across all initial angles. (**C-D**) Input noise correlation determines convergence angle. Simulations comparing uncorrelated versus temporally correlated Gaussian input noise (5% of input variance). (C) With uncorrelated noise, all odor pairs converge to complete orthogonality (90°) regardless of starting angle. (D) With temporally correlated noise, pairs converge to approximately 60°, matching experimental observations. Eight different starting angles tested (15°-85°). Final angles measured as average of last 50 trials.

**Supplementary Figure 4.**
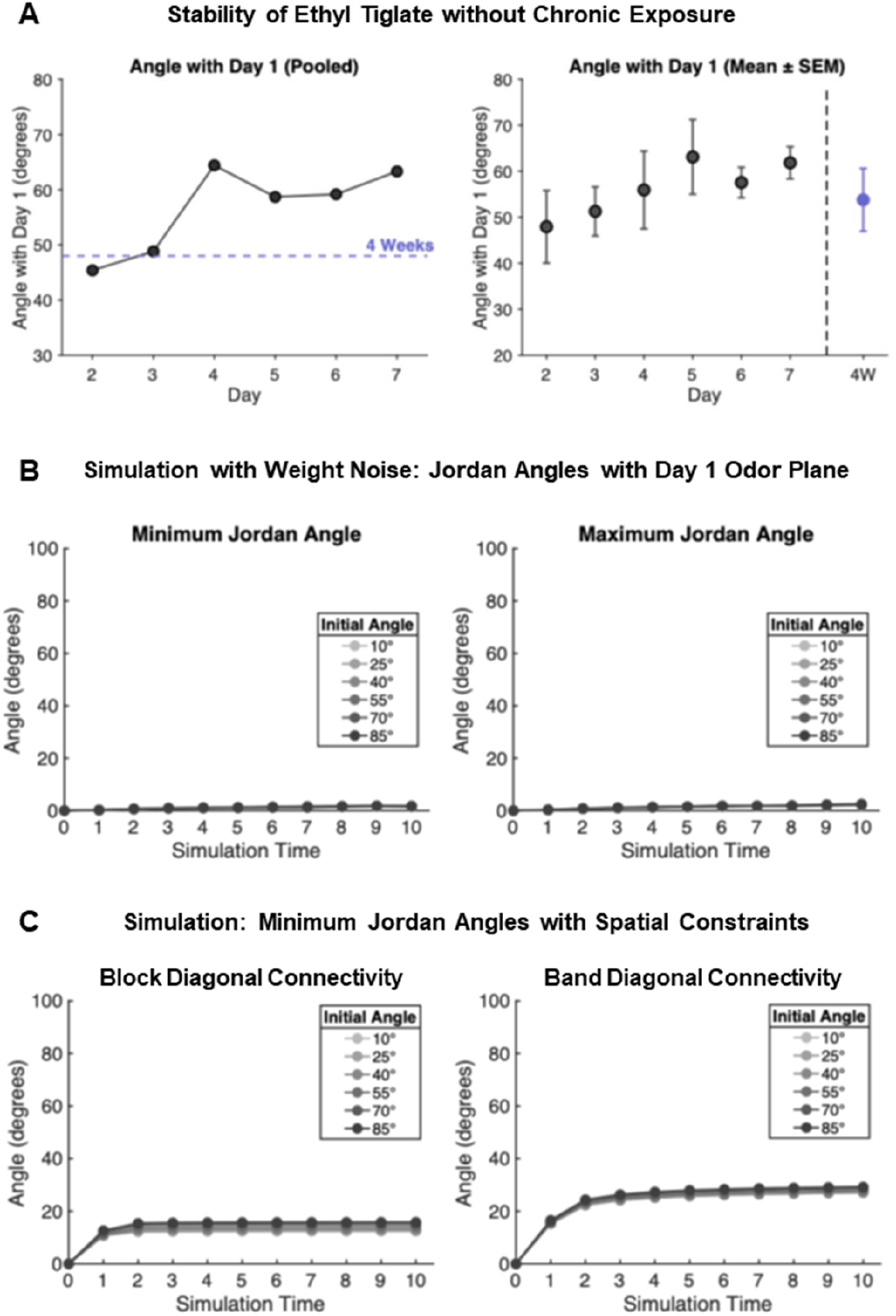
Representational drift requires chronic exposure and emerges from structural connectivity constraints. **(A) Drift magnitude from chronic exposures exceeds baseline variability from minimal exposure**. Angular change of Ethyl Tiglate (referred to as odor 2) representations relative to Day 1, compared with data from an independent study that also exposed animals to this odor at 4-week intervals with no intervening exposure (Shani-Narkiss et al., 2023; Shani-Narkiss et al., 2020). Left: pooled population vectors (N = 217 neurons, 6 mice for chronic exposure; N = 294 neurons, 7 mice for 4-week study). Right: daily angles of odor representation vectors of individual animals to Day 1 (Mean ± SEM). The comparison shows that chronic exposure produces substantially larger angles to initial exposure than the 4-week baseline, demonstrating that drift is exposure-dependent rather than spontaneous. **(B)** Weight update noise alone does not produce subspace rotation. Simulation results for fully connected MC-GC networks with random normal weight noise (magnitude comparable to learning-driven changes). No Jordan angle rotation is observed. **(C)** Structured connectivity produces drift in both minimum and maximum Jordan angles. Minimum Jordan angles for block-diagonal (left) and band-diagonal (right) connectivity patterns, using identical parameters as Fig. 4G and 4J. Both connectivity structures produce changes in minimum Jordan angles, demonstrating that the drift phenomenon affects multiple dimensions of the representational subspace.

**Supplementary Figure 5.**
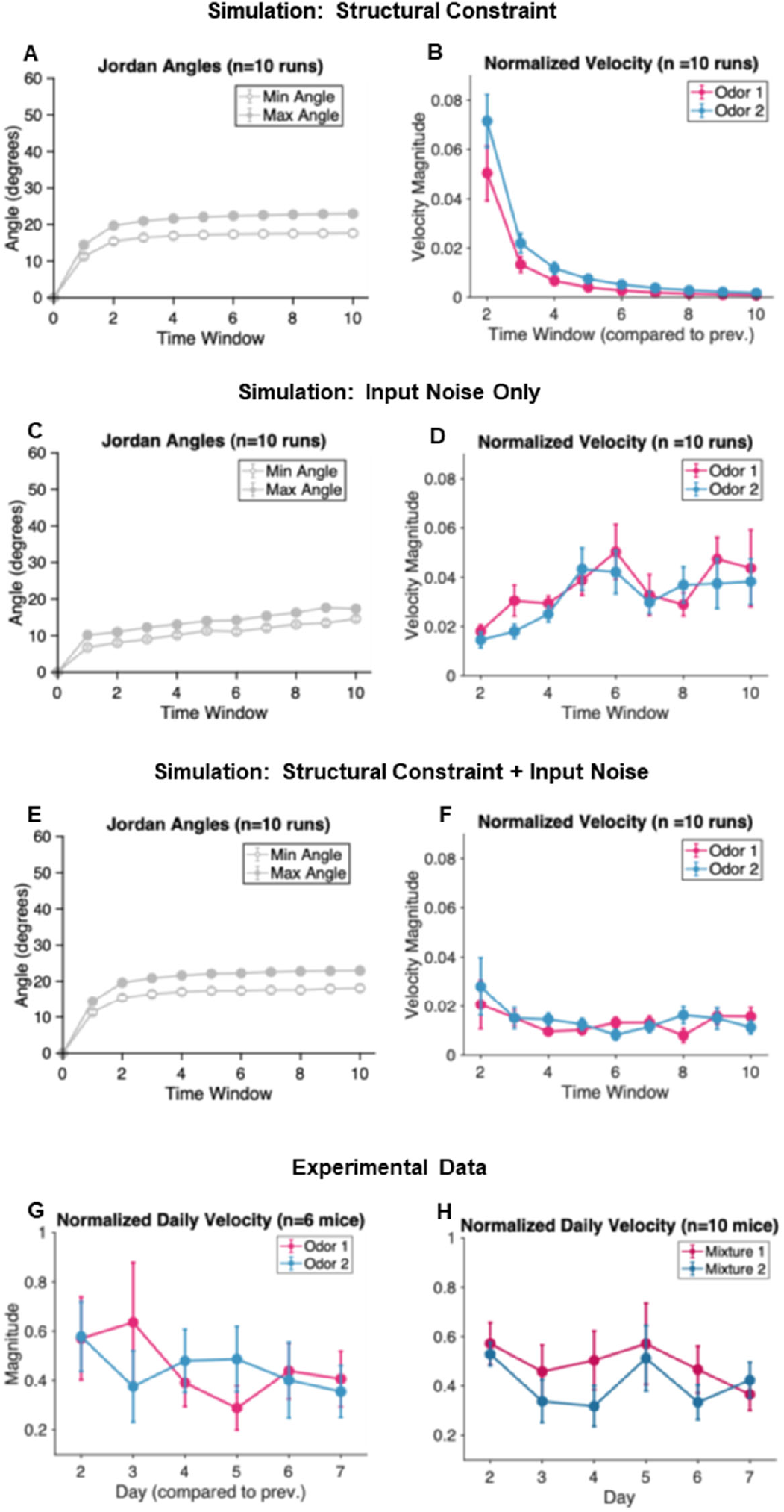
Distinguishing contributions of structural connectivity constraints versus input noise to representational drift. (**A-F**) Simulation analysis of drift mechanisms using normalized odor velocity. To isolate odor-specific drift effects from global subspace changes, we analyzed normalized velocity, the magnitude of change between consecutive time windows (normalized to remove decay effects from gain adaptation). Simulations used identical parameters except for the varied factors. Trial averages were computed by averaging over 100-trial windows; n=10 runs with different random seeds. Simulation data show mean ± SEM over random seeds. (A-B) Structural constraint alone. Block-diagonal connectivity produces substantial Jordan angle drift (A) with high initial velocity that rapidly decreases (B). (C-D) Input noise alone. Input noise (10% variance) produces modest Jordan angle changes (C) but increasing velocity over time (D), as noise overwhelms diminishing residual changes due to input dependent plasticity. (E-F) Combined structural constraint and input noise. Both factors together produce Jordan angle drift (E) with velocity dynamics (F) that show initial higher velocity followed by gradual decrease. **(G-H)** Experimental validation. Normalized daily velocity from experimental data shows patterns most consistent with combined mechanisms. (G) Dissimilar odor pairs. (H) Similar odor pairs. Data show mean ± SEM across animals.

**Supplementary Figure 6.**
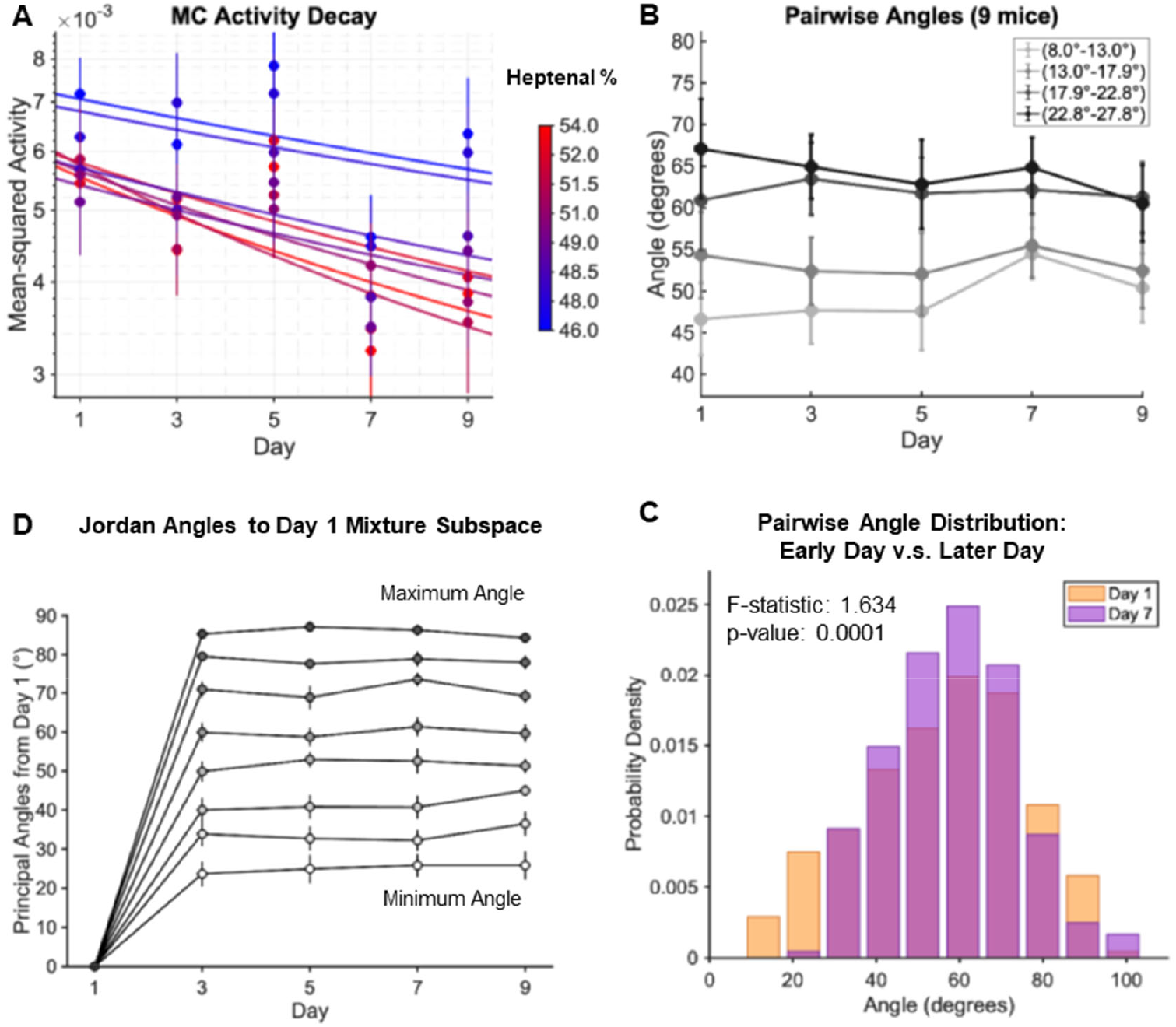
Neural representations for binary mixture panel exhibits similar representational changes as odor pairs during passive exposure. **(A)** Mean-field decay model fits for individual mixture responses. Mean-squared population activity (±SEM) across all mice for each of the 8 binary mixtures, color-coded by Heptanal percentage. Solid lines show best-fit power-law decay functions from our mean-field model, demonstrating that gain adaptation occurs consistently across the concentration gradient. **(B)** Pairwise angle changes depend on initial separation. Angular changes between mixture representations, binned by initial Day 1 angles into 4 equally spaced intervals (containing 4, 7, 8, and 9 pairs respectively). Data show mean ± SEM across animals. Pairs with larger initial angles tend to decrease (convergence), while pairs with smaller initial angles tend to increase (separation), consistent with convergence toward an intermediate angle. **(C)** Distribution of pairwise angles narrows over time. Histogram comparing all pairwise angles between the 8 mixtures on Day 1 (orange) versus Day 7 (purple) across all animals (28 pairs × 9 animals = 252 data points). The distribution becomes more concentrated around 50-60°, supporting the theoretical prediction of angle convergence. F-statistic and p-value from two-sample F-test for equal variances. **(D)** Jordan angles quantify subspace drift for mixture representations.

## Supplemental Information

### 1 Learning in the GC-MC circuit

Our model is based on the following equations describing the responses of MCs *m* ∈ ℝ^*M*^, and responses of GCs *a* ∈ ℝ^*N*^ :

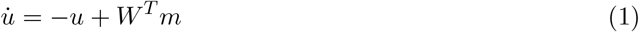

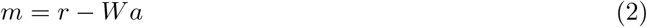

with

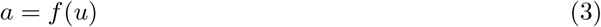

where *u* ∈ ℝ^*N*^ is the membrane voltage of the GCs. Here, *W* ∈ ℝ^*M* ×*N*^ is the connectivity matrix from GCs to MCs, mediating local inhibitory feedback, *r* is input from the olfactory sensory neurons (OSNs). Since the connection is via dendrodendritic synapses, excitation from MCs to GCs occurs through *W* ^*T*^, and we assume equal synaptic strengths for simplicity. For tractability, we consider linear activation *f* (*u*) = *θu, θ* being a shared scalar parameter for all GCs, and we assume instantaneous MC responses.

The differential equations (2)-(3) admit a Lyapunov function:

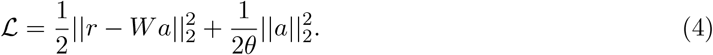

as the temporal behavior of the system can be viewed as gradient descent, i.e., 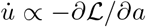, and the time derivative of the Lyapunov function is always negative:

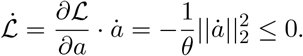

Inspecting the Lyapunov function, the first term describes the *error* in the representation of the inputs *r* by the GCs, and minimization of this difference leads to the most accurate approximation of the receptor neuron activity by the GCs. The second term of the Lyapunov function represents a regularization cost imposed on the GC firing, which can be seen as a constraint favoring small activation magnitudes. Therefore, the dynamical system can be thought of as performing regularized linear reconstruction, where the GCs learn a representation *a* that reconstructs the input *r* using the fixed basis *W* while penalizing large activations, and the output *m* to the cortex is the residual error between the input and its reconstruction.

The same Lyapunov function also provides a natural framework for synaptic plasticity. Taking the gradient of ℒ with respect to the connectivity matrix *W* yields:

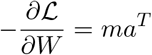

This gradient gives rise to the classic Hebbian learning rule, where synaptic strength increases when both pre-synaptic GC activity (*a*) and post-synaptic MC activity (*m*) are simultaneously active.

Remarkably, this learning rule emerges naturally from the same optimization principle that governs neural dynamics, suggesting that both inference (finding appropriate representations) and learning (adapting synaptic weights) can be understood as minimizing the same objective function. The resulting plasticity rule is both i) local, requiring only information available at individual synapses, and ii) unsupervised, making it biologically plausible while providing a principled approach to dictionary learning in the GC-MC circuit.

#### 1.1 Separation of Time Scale

We study this system assuming fast dynamics in the bulb and slow synaptic plasticity, which occurs when the animal is repeatedly exposed to odor inputs.

First, with every inhalation, the dynamics quickly reach equilibrium, and *a*^∗^ and *m*^∗^ can be solved to take the form:

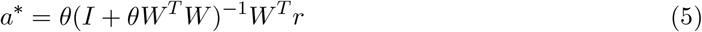

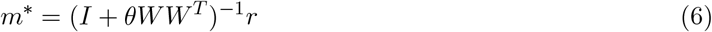

Then, the synaptic weights update incrementally according to Δ*W* = *η m*^∗^*a*^∗*T*^ − *λW*, where *η* is learning rate, and *λ* is weight decay (turnover).

Note that we do not constrain the sign of *W* ‘s elements. While biological synapses are non-negative, our analysis focuses on the mathematical structure where *WW* ^*T*^ symmetry is preserved, which holds regardless of *W* ‘s positivity constraint.

### 2 Mean-field Decay

We consider a simplified network model to understand the decay of mitral cell population activity over learning. The model consists of *M* MCs with input vector **r** = [*r*_1_, *r*_2_, …, *r*_*M*_]^*T*^. The MCs are connected to a single granule cell to capture the inhibitory dynamics while maintaining analytical tractability. We used vector bold font in this section for clarity of vectorized analysis.

The simplified network is governed by the following equations:

1. **GC activities:** 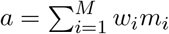
2. **MC activities:** *m*_*i*_ = *r*_*i*_ − *w*_*i*_*a*
3. **Hebbian weight update:** 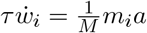

where *τ* is the synaptic time constant, and *w*_*i*_ are the dendrodendritic synaptic weights. Scaling the learning rate by 1*/M* ensures that larger populations do not learn faster just because they have more neurons contributing to the weight updates, and the normalization maintains consistent dynamics across population sizes.

Substituting the GC equation into the MC equation:

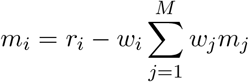

We can solve for the self-consistent solution for *a* by summing over *i* for *m*_*i*_ = *r*_*i*_ − *w*_*i*_*a* and get:

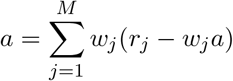

Solving it for *a* we obtain:

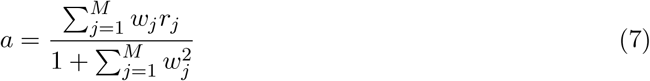

#### Weight dynamics decomposition

We can decompose the weight vector as:

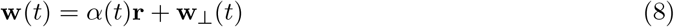

where 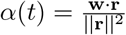 is the projection coefficient onto the input direction, and **w**_⊥_· **r** = 0. We evolute the evolution fo the parallel and perpendicular components next. The goal is to show that the orthogonal direction would diminish over time, and then weight dynamics, and MC activity can be expressed using scalar dimension *α*(*t*).

#### Parallel component evolution

Since 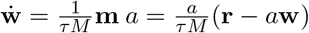,

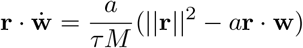

With orthogonality constraint, we also have:

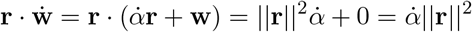

With the above two equations for 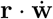, we can readily derive:

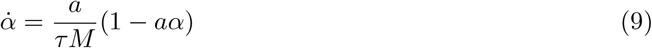

#### Perpendicular component evolution

From 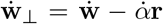 and the derived dynamics for 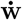 and 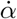:

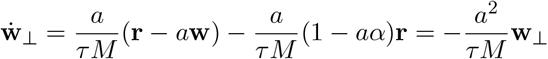

Since −*a*^2^*/τ <* 0, the perpendicular component decays near exponentially:

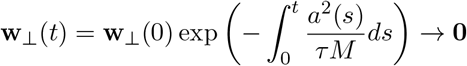

After the perpendicular component decays, **w**(*t*) ≈ *α*(*t*)**r**, which simplifies the collective variables:

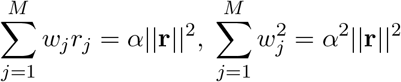

and therefore we can write *a* in terms of *α* and **r**:

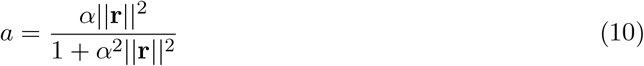

The evolution 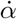 becomes:

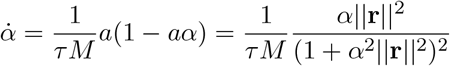

For *α*^2^||**r**||^2^ ≫ 1, which is equivalent to 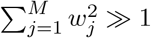 (valid when the synaptic weights have grown substantially large),

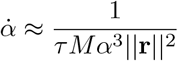

This approximation captures the asymptotic regime where the weights have strengthened considerably through Hebbian learning.

This separable equation integrates to:

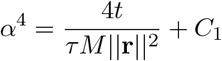

On the other hand, we can derive for individual MC activity with expression of *α* too:

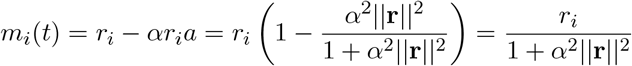

Therefore, population second moment:

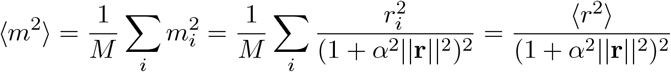

Substituting expression for *α* into the second moment expression and using ||**r**||^2^ = *M* ⟨*r*^2^⟩:

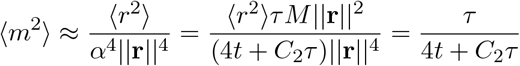

This provides an explicit functional form for fitting experimental data, with parameter *τ* representing the effective synaptic time constant, and *C*_2_ a shifting constant. In the main text, we use *C* = *C*_2_ to denote this constant, and therefore we have:

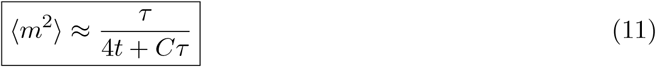

The mean-field approach gives us the inverse-time decay, which provides mechanistic insight for how weight alignment with input patterns drives the reductions of mitral cell population activities.

### 3 Learning on Single Odor Input

#### 3.1 Residual *m* Aligns to *r*

In this section, we show that the residual for a single input during learning with Hebbian plasticity aligns with the input. By “alignment,” we mean that *m* is parallel to *r*, achieving cos(*ψ*) = 1 where *ψ* is the angle between *m* and *r*. We assume the network has been pre-trained on samples drawn from the same distribution as the input, a centered Gaussian distribution. This initialization controls for distributional novelty effects, ensuring that observed alignment dynamics reflect learning of the specific input pattern *r* rather than adaptation to an unfamiliar input statistics.

**Setup:** Before exposing the network to a specific input *r*, we assume it learns from samples *r*′ ~ 𝒩 (0, *σI*) in dimension *M*.

**Lemma 1:** After pre-training on centered inputs, the residual *m* aligns with any new input *r* at first exposure.

*Proof*. During pre-training, the network learns from samples *r*′ ~ 𝒩 (0, *σ*^2^*I*). In the limit of many samples, the sample covariance becomes:

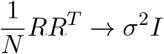

At steady state with weight decay *λ*, the dynamics satisfy:

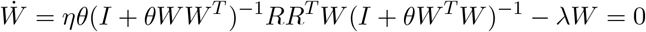

With SVD *W* = *USV* ^*T*^, this becomes:

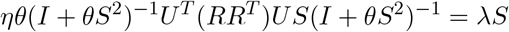

Since *U* ^*T*^ (*σ*^2^*I*)*U* = *σ*^2^*I*, we get:

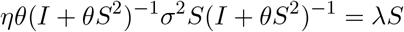

For each singular value *s*_*i*_:

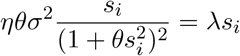

Solving for *s*_*i*_:

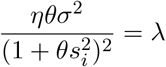

This gives the same solution for all *i*:

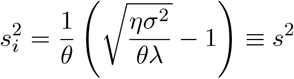

For the pre-trained network, all singular values are equal: *S* = *sI*. The residual for any new input *r* is:

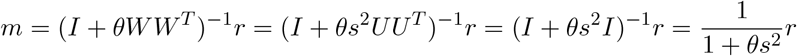

Therefore, *m* is perfectly aligned with *r* during first exposure.□

**Lemma 2:** After *W* update on *r* after first exposure, *m* aligns with *r*, and same applies to all future updates.

*Proof*. We call the weight matrix after pretraining *W*_0_, with 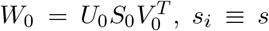, and that *W*_1_ = *W*_0_ + *η ma*^*T*^.

Therefore *W*_1_ can be written as:

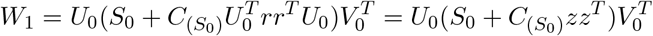

where 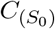 is diagonal with all entries of same values, and we also simplify by defining 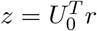.

Let 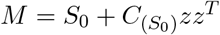. Since *S*_0_ = *sI* and 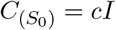 for some scalar *c*, we have:

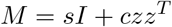

The eigenvalues of *M* are: *λ*_1_ = *s* + *c*∥*z*∥^2^ with eigenvector 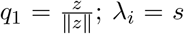 for *i* = 2, …, *M* with eigenvectors orthogonal to *z*.

Let *M* = *Q*Λ*Q*^*T*^,

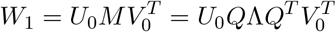

Then we know singular vectors *u*_*i*_ of *W*_1_, *u*_*i*_ = *U*_0_*q*_*i*_. Since 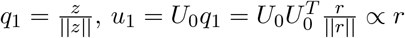.

Now, for the residual with updated weights, since 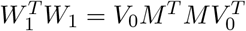, they share singular values:

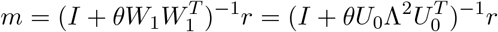

Since *r* ∝ *u*_1_, we can write *r* = *αu*_1_ for some scalar *α >* 0. Then:

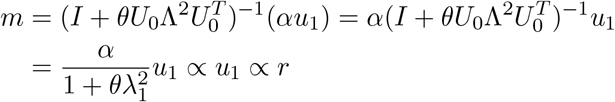

Therefore, *m* remains aligned with *r* after the first weight update.

##### Inductive argument

By induction, this alignment is preserved for all future updates. At each exposure to *r*, the Hebbian update Δ*W* = *ηma*^*T*^ further strengthens the connection in the *r* direction since both *m* and *a* are aligned with *r*. This increases the singular value *λ*_1_ corresponding to the *r* direction while leaving other singular values unchanged. Since the leading eigenvector *q*_1_ remains proportional to 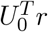 (and hence *u*_1_ ∝ *r*), the alignment is preserved through all subsequent updates.□

### 3.2 Presence of Weight Noise

In this section we show that there is no systematic rotation in expectation with stochasticity in weight update. For systematic rotation to exist in the expected trajectory, we would need the infinitesimal expected change in *m, δm*, to contain a part ∝ *Jm*, where *J* is an antisymmetric rotation matrix. Our goal is to inspect 𝔼[*δm*] = 𝔼[*m*(*t* + 1)] − 𝔼[*m*(*t*)] when *δm* becomes very small.

Recall that we have arrived at a succinct form of *m*, expressing it with *W* and *r*:

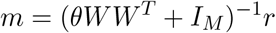

Let’s analyze how a single noisy update affects *m*. At time step *t*:

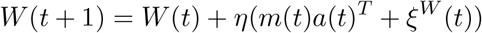

where *ξ*^*W*^ (*t*) has i.i.d. entries with variance *σ*^2^ ≤ *η*^2^||*ma*^*T*^ ||^2^.

Some key properties of the independent noise *ξ*^*W*^ :

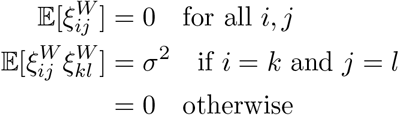

Let’s write the update as a perturbation:

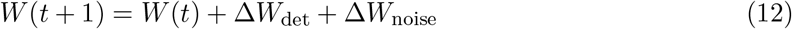

where:

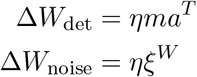

Using the relationship for *m*(*t* + 1):

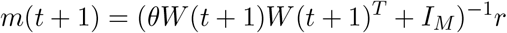

Using the expression of eq (12), multiply out (*W* + Δ*W*_det_ + Δ*W*_noise_)(*W* + Δ*W*_det_ + Δ*W*_noise_)^*T*^, we get:

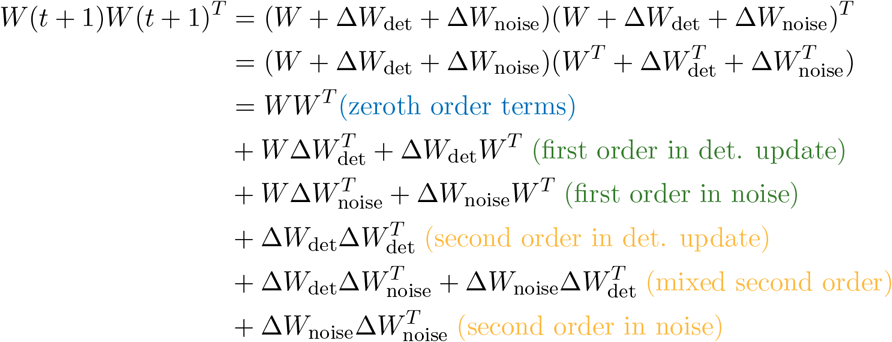

Now, substituting the Δ*W*_det_ = *ηma*^*T*^ and Δ*W*_noise_ = *ηξ*^*W*^ back, we get:

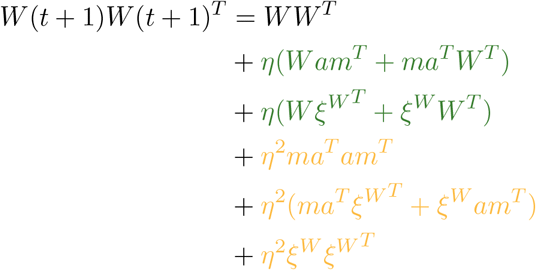

#### First-order noise terms

When we multiply *W* (*t* + 1)*W* (*t* + 1)^*T*^, the first-order terms in noise are:

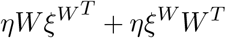

For each entry (*i, j*) in this matrix:

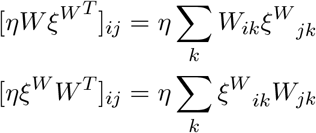

Therefore:

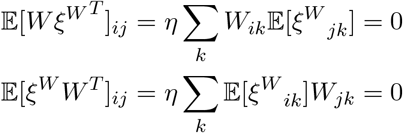

This shows that first-order noise terms vanish in expectation, and therefore any directional changes from noise must come from higher-order terms. This helps explain why when noise is comparable to the input (*σ* ~ *η*||*ma*^*T*^ ||) has minimal effect on the direction of *m*.

#### Second-order terms

The second-order terms in *W* (*t* + 1)*W* (*t* + 1)^*T*^ are:

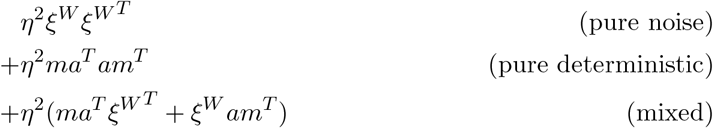

Taking expectations, same as in the pretraining session:

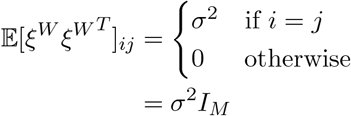

The mixed terms vanish in expectation since 𝔼[*ξ*^*W*^] = 0.

#### 3.2.1 Approximating 𝔼[*δm*]

Recall that *m* = (*θWW* ^*T*^ + *I*_*M*_)^−1^*r*. For the current state, we leave out the time subscripts and let’s write:

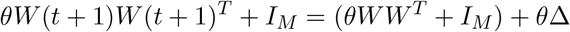

where Δ includes all update terms expanding *W* (*t* + 1)*W* (*t* + 1)^*T*^. Taking the expectation, then many terms zeroed out, and we are left with:

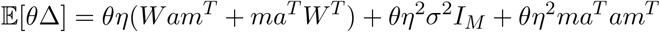

Now, treat *θWW* ^*T*^ + *I*_*M*_ as *A*, and the aggregated perturbation term *θ*Δ as *ϵB*, especially identify *η* with *ϵ*, we can use the below expansion to approximate (*θW* (*t* + 1)*W* (*t* + 1)^*T*^ + *I*_*M*_)^−1^:

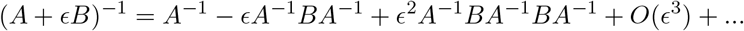

Applying this to our case, keeping only first order term,

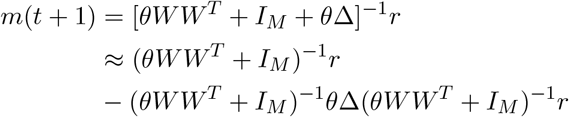

Taking expectation and use *m*(*t*) = (*θWW* ^*T*^ + *I*_*M*_)^−1^*r*

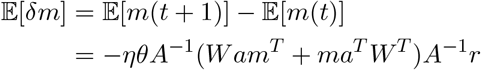

Note that *A*^−1^ is symmetric, and *Wam*^*T*^ + *ma*^*T*^ *W* ^*T*^ is a matrix summed with its transpose so always symmetric, thus there is no antisymmetric component. We conclude that when update *W* with isotropic Gaussian noise, there is no systematic rotation being introduced.

### 3.3 Presence of Input Noise

Assume that input is 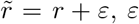, *ε* being random Gaussian with zero mean and a small variance, compared to the input variability. We find the expected converged 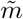 ‘s expression will give us information about the deviation from *r*’s direction.

Consider training with noisy input 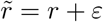, where *ε* ~ 𝒩 (0, *σ*^2^*I*). The residual becomes:

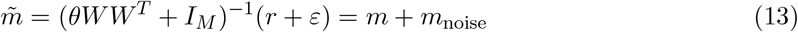

where *m* = (*θWW* ^*T*^ + *I*_*M*_)^−1^*r* and *m*_noise_ = (*θWW* ^*T*^ + *I*_*M*_)^−1^*ε*.

#### Steady-state analysis with input noise

The weight update becomes:

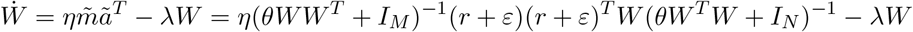

At steady state, 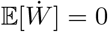. Since 𝔼[*ε*] = 0 and 𝔼[*εε*^*T*^] = *σ*^2^*I*:

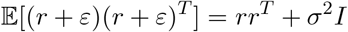

Using our standard SVD approach *W* = *USV* ^*T*^, the steady-state condition becomes:

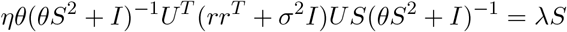

For this to be diagonal, we need *U* ^*T*^ (*rr*^*T*^ + *σ*^2^*I*)*U* to be diagonal. This requires *U* ^*T*^ *r* to have only one non-zero component, giving:

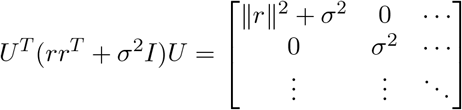

Therefore, 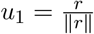 remains aligned with *r*, and the singular values satisfy:

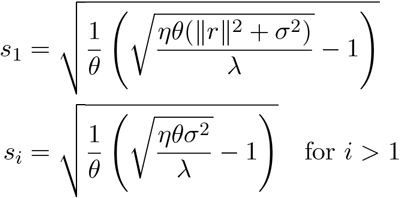

The residual thus decomposes as:

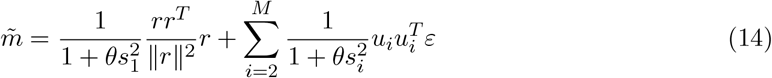

which shows that, the deterministic component *m* ∝ *r* maintains alignment with the input, but noise is filtered differently across directions: strongly suppressed along *r* (due to large *s*_1_), less suppressed in orthogonal directions (due to smaller *s*_*i*_). The noise magnitude is larger in directions orthogonal to *r*, creating bounded deviations from perfect alignment.

Moreover, with continued learning, the deterministic component of 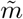 decays faster than the noise component. Since 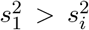 for *i >* 1, the deterministic term 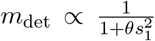 is suppressed more strongly than the noise terms 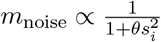. This leads to random walk-like drift in the orthogonal subspace, even though 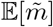 remains aligned with *r* in expectation.

### 3.4 Presence of Structural Connectivity

In realistic neural circuits, synaptic connectivity is constrained by anatomy. While analyzing arbitrary sparse connectivity matrices is intractable, we can gain insight by studying simplified cases that capture key features of olfactory bulb organization.

The olfactory bulb exhibits structured connectivity: MCs and GCs are arranged in layers with dense local connections due to physical proximity and limited dendritic spread, supplemented by sparse long-range connections. To understand how such connectivity patterns affect learning dynamics, we analyze two tractable cases that preserve essential structural features:

**1) Block diagonal connectivity:** *A* is block diagonal with all-to-all connections within blocks, modeling distinct local circuits.

**2) Banded connectivity:** *A* has a band matrix structure, modeling distance-dependent connection probabilities.

Both cases constrain the weight updates *W* ← *W* + *η*(*ma*^*T*^ ⊙ *A*), where ⊙ denotes element-wise multiplication, fundamentally altering the learning dynamics from the all-to-all case.

#### 3.4.1 Block-diagonal Connectivity

##### Intuition for the block diagonal case

consider *A* with two blocks:

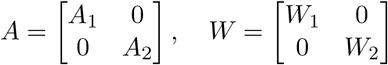

The key insight is that (*θWW* ^*T*^ + *I*_*M*_)^−1^ inherits the block structure:

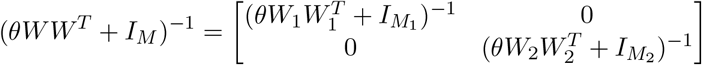

Therefore, for input 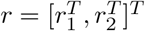:

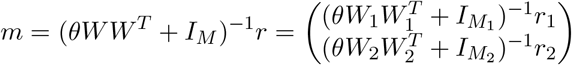

Key consequences for block constraint is that each block processes its input independently, and each block’s suppression is scaled by its own input, and this differential suppression by block creates an angle between the overall *m* and *r*.

For constant input *r* and block sizes that are approximately Gaussian-distributed *x* ~ 𝒩 (*µ, σ*^2^) (which approximates the binomial distribution under the assumption of large *M* and many blocks) with *K* = *M/µ* total blocks, the angle between *m* and *r* is:

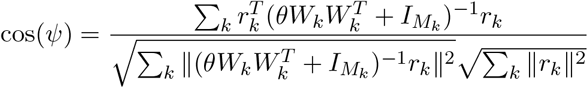

##### Steady state angle for varying block sizes

At steady state with weight decay *λ*, each block *k* satisfies:

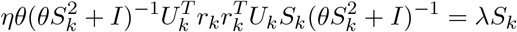

Following our standard analysis, the first singular vector *u*_*k*,1_ aligns with *r*_*k*_, giving:

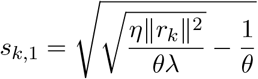

For each block, the residual becomes:

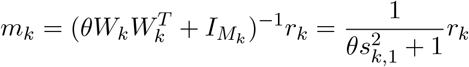

The overall angle between *m* and *r* is:

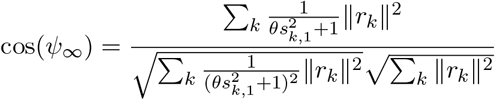

For constant input density, 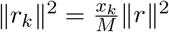 where *x*_*k*_ is the size of block *k*. Defining 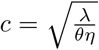 and substituting:

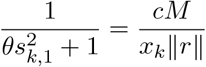

This yields:

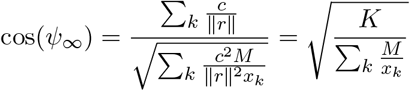

where *K* is the number of blocks.

##### Expected angle with block size variance

For *x*_*k*_ ~ 𝒩 (*µ, σ*^2^), using the approximation 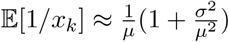:

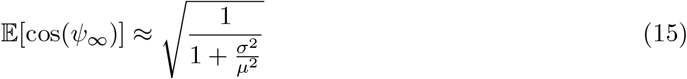

If we have uniform blocks (*σ* = 0), we would have perfect alignment (cos(*ψ*_∞_) = 1); on the other hand, if we have high variance,(*σ*^2^*/µ*^2^ ≫ 1): alignment would be poor (cos(*ψ*_∞_) → 0).

##### Analysis of input distribution and final angle

Now we fix the block size *d* with *K* total blocks, and analyze how input distribution affects alignment.

From our steady-state analysis:

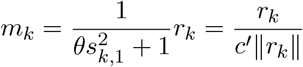

where 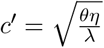.

The cosine of the angle between *m* and *r* becomes:

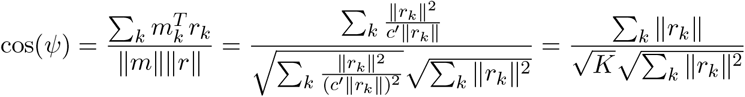

##### Gaussian input analysis

For each component *r*_*k,i*_ ~ 𝒩 (*µ, σ*^2^) independently, we have:

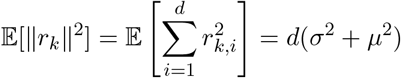

For the expected norm 𝔼[∥*r*_*k*_∥], since ∥*r*_*k*_∥^2^ follows a non-central chi-squared distribution, we can compute numerically:

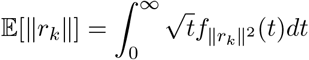

Therefore:

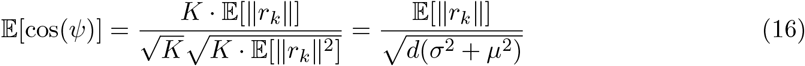

When we have uniform inputs (all ∥*r*_*k*_∥ equal), cos(*ψ*) = 1. When not, alignment depends on both mean *µ* and variance *σ* of inputs. The larger block count *K*, the smaller the block-wise variance, the greater the alignment (simulation see Fig. 4H in main).

### 3.4.2 Band-diagonal Connectivity

#### Intuition with band structure

consider connectivity matrix *A* with fixed bandwidth *k*:

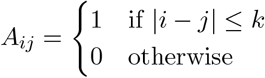

This constrains *W* to have the same band structure: *W*_*ij*_ = 0 if |*i* − *j*| *> k*.

Let’s inspect the band structure effects. For *B* = *θW* ^*T*^ *W* + *I*_*N*_, we can use the geometric series expansion:

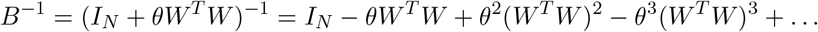

With power *n*, (*W* ^*T*^ *W*)^*n*^ has bandwidth 2*nk*, but with decreasing magnitude due to the *θ*^*n*^ factor (assuming *θ <* 1). The residual becomes:

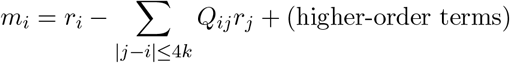

This shows that each *m*_*i*_ is primarily influenced by *r*_*j*_ within distance ~ 4*k*, creating a “sliding window” like effect, where adjacent regions share input components, therefore still creating local suppression patterns, similar to the block diagonal case.

## 4 Learning on Odor Pair

### 4.1 The Decorrelation Motif

**Theorem:** When the system learns two input vectors *r*_1_ and *r*_2_, their corresponding residuals *m*_1_ and *m*_2_ will be orthogonalized at steady state.

*Proof*. We will show that if *r*_1_ and *r*_2_ initially form angle *ϕ*, then *m*_1_ and *m*_2_ will each move away from the other input by angle 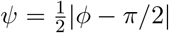, achieving orthogonality when *ϕ* ≠ *π/*2.

#### Coordinate system and eigendecomposition

Without loss of generality, consider two input vectors *r*_1_ and *r*_2_ with equal magnitude *α*. We choose coordinates such that:

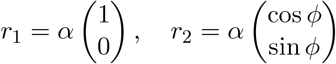

where *ϕ* is the angle between *r*_1_ and *r*_2_.

The covariance matrix becomes:

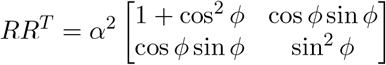

Solving the characteristic equation det(*RR*^*T*^ − *δI*) = 0 yields eigenvalues:

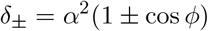

with corresponding normalized eigenvectors:

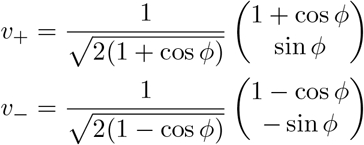

#### Steady-state weight matrix

At steady state with weight decay *λ*:

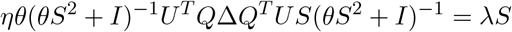

For this equation to hold with diagonal right-hand side, we require *Q* = *U*. Therefore, the left singular vectors of the steady-state *W* are *U* = [*v*_+_|*v*_−_]:

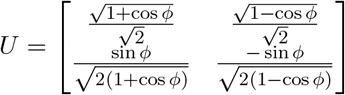

with singular values:

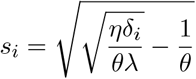

#### Residual computation and angle derivation

The residual for *r*_1_ is:

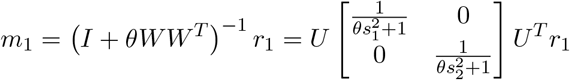

Defining 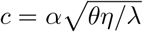 and substituting the expressions for *s*_*i*_ and *U*, we obtain:

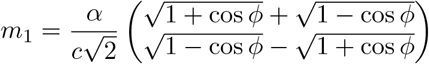

The angle *ψ* between *m*_1_ and *r*_1_ satisfies:

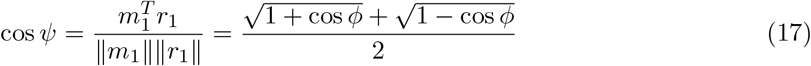

Now we show that this implies 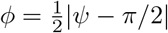:

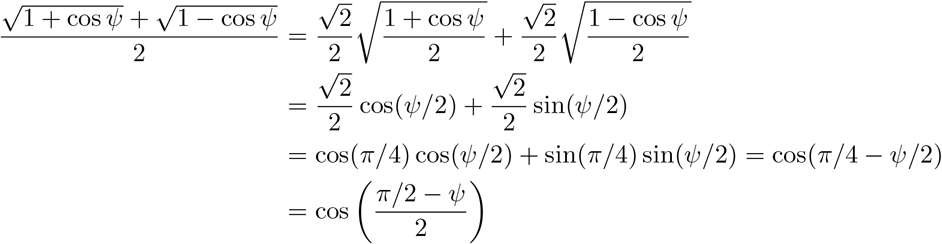

Since 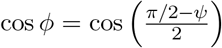, we conclude that 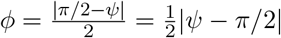.

#### Prediction for more than two inputs

Our prediction is that for *n* input odors, the system will achieve pairwise orthogonalization of all residuals *m*^*i*^. Each residual will move in the direction that maximizes its distance from the sub-space spanned by all other inputs, ultimately converging to a configuration where all residuals are mutually orthogonal. Complete orthogonalization can be achieved as long as the input dimension satisfies *M* ≥ *n*.

#### Prediction for learning input sets with lower rank

Consider a set of *n*_1_ input mixtures composed of *n*_2_ underlying odorants, where *n*_2_ *< n*_1_. Although there are *n*_1_ distinct inputs, the covariance matrix *RR*^*T*^ has rank *n*_2_, constraining all inputs to lie within an *n*_2_-dimensional subspace. We predict that the system will learn this covariance structure and optimally separate the inputs within the subspace they span. The residuals will achieve maximal separation subject to the constraint that they remain orthogonal to the null space of the input covariance.

### 4.2 Temporally correlated noise

In Section 3.3, we showed that with uncorrelated input noise, the residual *m* aligns with *r* in expectation for a single input. Therefore, with two inputs and independent noise, *m*_1_ and *m*_2_ will follow their expected decorrelation trajectory as derived in the noiseless case.

However, motivated by trial-to-trial noise correlations observed in neural responses, we now consider correlated noise between inputs: 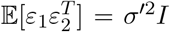 where *σ*′ *< σ*. These shared fluctuations across trials prevent perfect decorrelation.

For two noisy inputs 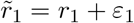 and 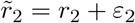, we analyze the residual correlation:

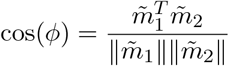

The numerator:

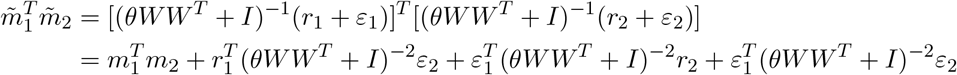

Taking expectations and using 𝔼[*ε*_*i*_] = 0:

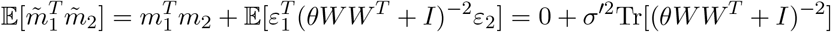

where the first term vanishes at convergence due to perfect decorrelation in the noiseless case. For denominator similarly:

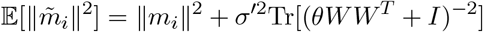

Defining *T* = Tr[(*θWW* ^*T*^ + *I*)^−2^]:

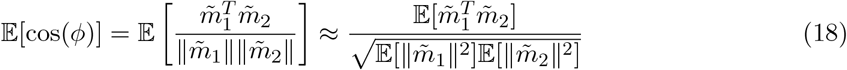

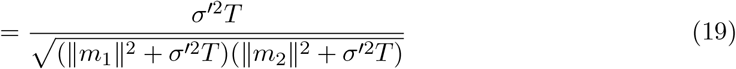

Inspecting this equation, we can see that perfect decorrelation is impossible: with correlated noise (*σ*′ *>* 0), 𝔼[cos(*ϕ*)] *>* 0, residuals cannot achieve perfect orthogonality.

From our following two-input analysis, we can realitively easily show that ∥*m*_1_∥^2^ = ∥*m*_2_∥^2^ regardless of the initial angle *ϕ* between *r*_1_ and *r*_2_.

With *W* = *USV* ^*T*^, we have at steady state:

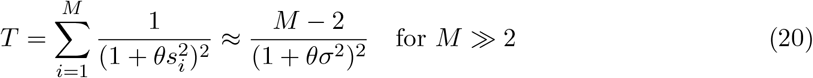

Since *T* is dominated by the noise subspace dimensions and is similar across all pairs, all odor pairs should converge to approximately the same residual correlation angle and the angle is controlled by mostly the noise magnitude.

This prediction is confirmed in stimulation (see Fig. 3G in main).

